# Tissue-specific tolerance mechanisms and lymph node co-drainage converge to shape T cell immunity in the upper digestive system and regulate pancreatic cancer progression

**DOI:** 10.64898/2025.12.02.691870

**Authors:** Yixuan D. Zhou, Peter Wang, Emily Schaffer, Macy R. Komnick, Hailey Brown, Gwen M. Taylor, Kay L. Fiske, Colin Sheehan, Terence S. Dermody, Alexander Muir, Daria Esterházy

## Abstract

The liver, pancreas, and duodenum share lymph nodes (LNs), providing a unique system to examine how tissue origin of self-antigens shapes T cell fate. Comparing mice expressing ovalbumin (OVA) from distinct subcellular compartments, we found that cytosolic OVA from liver or pancreas, but not gut, was immunologically ignored. High-dose hepatic secreted OVA triggered antigen-specific T cell deletion, whereas secreted pancreatic and intestinal OVA induced regulatory T (Treg) cells, revealing immunological ignorance, clonal deletion and Treg cell generation as tissue-specific tolerance mechanisms. Of these, LN co-drainage only influenced Treg cell induction, establishing gut-pancreas-liver axes: Intestinal viral infection rendered hepatocyte- and exocrine pancreas-specific T cells inflammatory; liver injury promoted pancreas- and gut-directed responses. These self-reactive T cells caused tissue destruction but enhanced pancreatic tumor control when neoantigen OVA was secreted but not cytosolic. Thus, LN co-drainage and tissue-specific tolerance mechanisms jointly shape immune homeostasis and disease susceptibility in the upper digestive system.

## Introduction

The immune system must balance protective reactions against microbial pathogens and malignancies with tolerance to self and innocuous foreign antigens. Achieving this balance requires finely tuned responses tailored to the distinct environments of individual organs, underscoring the importance of compartmentalization in immunity. Immune cells acquire organ-specific transcriptional and functional profiles shaped by local cues such as antigen exposure, cytokines, and stromal signals^1,2^. Similarly, lymph nodes (LNs) draining different tissues provide specialized niches where adaptive immune responses are initiated in a context-dependent manner^3^. Immune compartmentalization exists even within a single organ, with the regionalization of gut draining LNs being a prime example^4^. LNs along the gut are immunologically distinct: in response to dietary antigens, proximal gut-draining LNs favor peripheral regulatory T (pTreg) cell induction, whereas distal LNs favor pro-inflammatory RORγt^+^ T cells. These differences parallel distinct dendritic cell profiles^4^ and stromal cell milieus^5^.

The strong influence of the gut LN milieu is further exemplified by LN sharing between organs. The duodenum, liver and pancreas share LNs^6^, and this anatomical arrangement biases pancreatic β-cell-specific T cells toward the Treg cell lineage under steady state but can redirect them towards pro-inflammatory lineages during duodenal viral infection^6^. In addition, we recently found that the subcellular origin of self-antigens originating from the gut can influence T cell fate within the same gut-draining LNs (gLNs)^7^. This effect is mediated by distinct dependencies on antigen-presenting cell subsets and translate into differential disease susceptibilities^6,7^. Since pancreatic and hepatic antigens are likely presented primarily by cDC1, the only migratory APC subset in these tissues^6^, it remains unclear whether the same rules governing antigen access and T cell fate in the gut also apply to antigens derived from these tissues. Understanding this is critical, as it will determine the circumstances under which the liver and pancreas are susceptible to T cell-mediated autoimmunity or cancer control, and when these responses can be shaped by LN co-drainage. This question is particularly relevant in the context of pancreatic ductal adenocarcinoma (PDAC), a malignancy with poor prognosis, frequent hepatic metastasis, and limited responsiveness to immunotherapies^8^.

LN sharing provides an excellent platform to interrogate whether T cell outcomes in response to self-antigens are primarily shaped by the tissue of origin or by the environment of the draining LN. In this study, we investigated whether the default peripheral tolerance mechanism of the gut, pTreg cell induction, also applies to self-antigens derived from the liver and pancreas when assessed within the same, shared LNs and using a standardized antigen, ovalbumin (OVA). We then examined how tolerance programs can be perturbed by selectively inducing inflammation either in the antigen-expressing tissue or in a co-drained organ. Finally, we assessed how tissue origin, subcellular localization and LN co-drainage intersect in dictating the susceptibility to T cell rewiring in the context of PDAC.

## Results

### Tissue-specific tolerance mechanism, subcellular source of self-antigen and antigen dose dictate T cell fate of self-reactive CD4^+^ T cells at homeostasis

To determine how the same self-antigen shapes T cell responses when expressed in different tissues within a shared lymphatic network, we engineered three mouse lines that express OVA in distinct forms, secreted (sOVA), cytosolic (cOVA), or transmembrane (tmOVA), under control of tissue-specific Cre drivers^7^. Using inducible *Alb^Cre-ERT2^* for hepatocytes, *Ptf1a^Cre-ERT2^* for exocrine pancreas and *Villin^Cre-ERT2^* for intestinal enterocytes, we expressed OVA in the liver, exocrine and endocrine pancreas, and gut^7^, respectively (Fig. S1A-D). Immunofluorescence imaging confirmed the subcellular distribution of each OVA variant (Fig. S1E), and no cross-expression was detected in non-target tissues within the triad (Fig. S1F-H).

To assess homeostatic CD4⁺ T cell responses, we adoptively transferred OT-II cells into OVA-expressing mice and quantified their proliferation in draining LNs 96 h after transfer. At steady state, tmOVA induced robust OT-II proliferation in liver and pancreas-draining LNs, whereas cOVA did not (Fig. 1A-D). The degree of OT-II proliferation mirrored the relative drainage of each tissue into the co-drained LNs (Fig. 1B, D) as previously identified^6^, and in line with the observation that tmOVA is seen by the immune system through ectodomain shedding^7^. A similar drainage hierarchy was observed for β-cell self-antigens, using *Ins1^Cre-ERT2^*, in which only sOVA and tmOVA elicited detectable proliferation (Fig. S1I). However, the number of OT-II cells recovered from each LN was insufficient to allow phenotypic analysis. To overcome this problem, we used streptozotocin (STZ) in the subsequent experiments to selectively induce β-cell death, enhancing antigen availability in draining LNs for phenotypic profiling. In contrast, all three OVA forms expressed in the gut elicited strong T cell proliferation^7^, consistent with the constant tissue turnover in the intestine, suggesting immune ignorance as the main mechanism for cytosolic self-antigen tolerance in tissues with low cell turnover. When hepatocyte or pancreatic acinar cell death was induced in mice expressing cOVA, OT-II cells proliferated in the respective draining LNs, reinforcing this notion (Fig. 1E, F).

**Figure 1.**
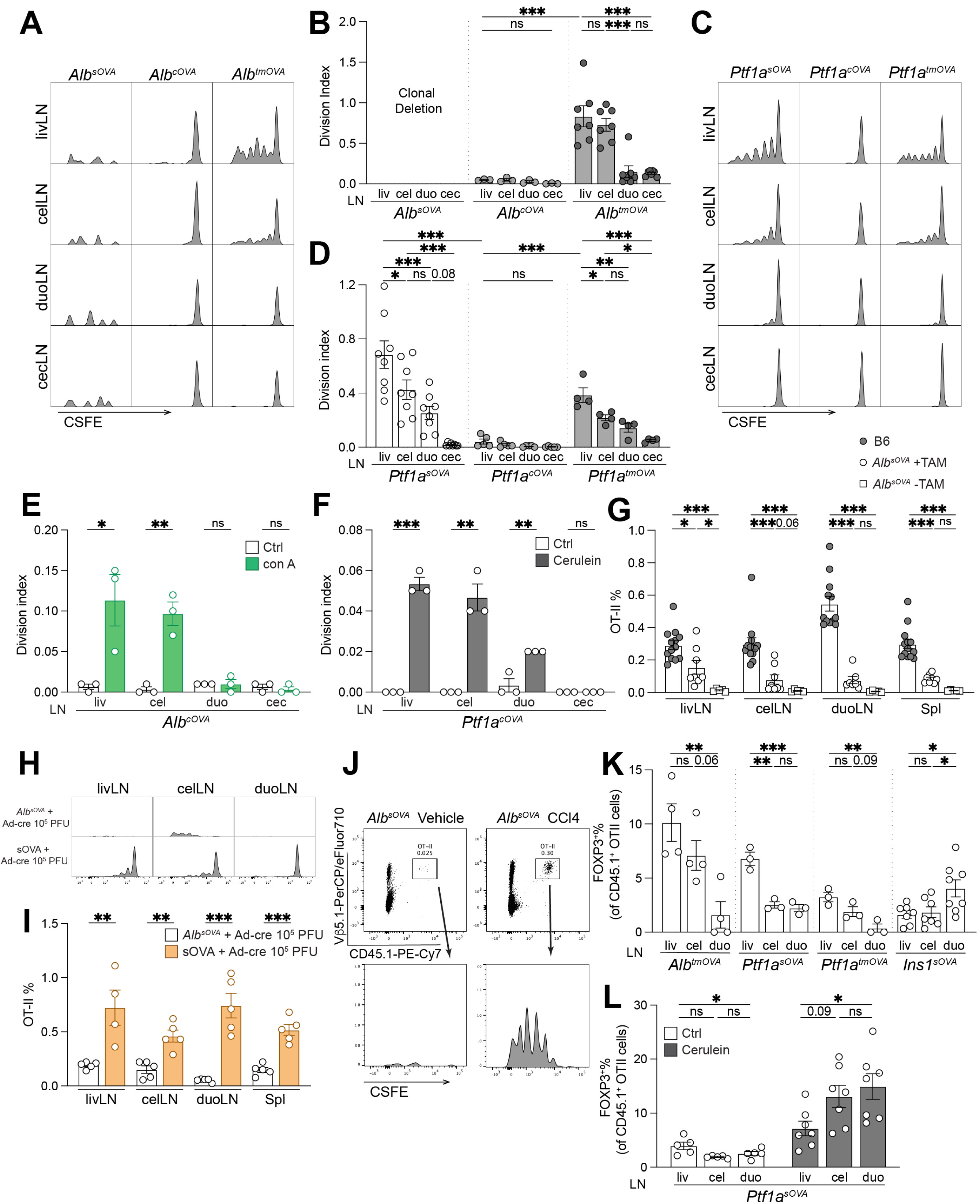
Tissue-specific tolerance mechanism, subcellular source of self-antigen and antigen dose dictate T cell fate of self-reactive CD4^+^ T cells at homeostasis. **A-B**. CFSE dilution flow plots (**A**) and division index (**B**) of OT-II cells in the indicated LNs 96 h after transfer into *Alb^sOVA^*, *Alb^cOVA^* or *Alb^tmOVA^* mice (*n* = 3-7 per group, as indicated by symbols). **C-D**. CFSE dilution flow plots (**C**) and division index (**D**) of OT-II cells in the indicated LNs 96 h after transfer into *Ptf1a^sOVA^*, *Ptf1a^cOVA^* or *Ptf1a^tmOVA^*mice (*n* = 4-8 per group, as indicated by symbols). **E**. Division index of OT-II cells in the indicated LNs 96 h after transfer into *Alb^cOVA^* mice treated with con A (*n* = 3 per group). **F**. Division index of OT-II cells in the indicated LNs 96 h after transfer into *Ptf1a^cOVA^* mice treated with cerulein (*n* = 3 per group). **G.** Frequencies of OT-II among total CD4^+^ T cells in the indicated LNs and spleen 96 h after transfer into OVA-naïve, tamoxifen-induced and non-induced *Alb^sOVA^* mice (*n* = 7-11 per group, as indicated by symbols). **H-I.** CFSE dilution flow plots (**H**) and frequencies of OT-II among total CD4^+^ T cells (**I**) in the indicated LNs of adeno-infected *Alb^sOVA^* and adeno-infected sOVA mice (*n* = 5 per group). **J.** Flow plots of OT-II cells in livLN of *Alb^sOVA^*mice treated with CCl_4_. **K**. Frequencies of FOXP3^+^ among total OT-II cells in the indicated LNs 96 h after transfer into *Alb^tmOVA^*, *Ptf1a^sOVA^*, *Ptf1a^tmOVA^* and *Ins1^sOVA^*mice (*n* = 3-8 per group, as indicated by symbols). **L**. Frequencies of FOXP3^+^ OT-II cells in the indicated LNs 96 h after transfer into *Ptf1a^sOVA^* mice treated with cerulein (*n* = 5-7 per group, as indicated by symbols). Data are representative of two independent experiments in B, D, E-G, K, L. * p<0.05, ** p<0.01, *** p<0.001 by one-way ANOVA and t-test.

Pancreatic sOVA led to stronger OT-II proliferation than tmOVA (Fig. 1C, D), in line with the observation that OVA released from tmOVA is present at lower levels than sOVA^7^. In contrast, hepatic sOVA, but not tmOVA, induced rapid OT-II clonal deletion (Fig. 1A, B). Since within the liver we observed that much more sOVA than tmOVA was expressed on the protein level (Fig. S1C), we speculated that hepatic antigen dose, rather than an inherent difference between secreted and transmembrane-shed antigen, was responsible for this striking difference in effect on CD4^+^ T cells. To test if antigen alone drove this deletion, we first attempted to lower OVA expression by reducing tamoxifen dosing. However, we discovered that *Alb^Cre-ERT2^*exhibits sufficient basal Cre activity even without tamoxifen to induce *OVA* expression (Fig. S1J, K). The expression level was comparable to that in tamoxifen-induced *Ptf1a^sOVA^* mice (Fig. S1K, L), which is a pertinent comparison since the liver and pancreas share similar LN drainage pattern, and was sufficient to delete OT-II cells (Fig. 1G). Such leakiness is not uncommon for tamoxifen-inducible Cre systems^9^, though not previously reported for *Alb^Cre-ERT2^*. In our system, high *Alb* expression and close loxP spacing likely both contribute to spontaneous recombination. We therefore delivered an adeno-Cre virus to the liver of Cre-negative sOVA mice by tail-vein injection instead. We chose an adenovirus dose that does not prevent OT-II cell deletion in *Alb^sOVA^* mice (Fig. 1H, I), as we discovered that high-dose viral infection can break this tolerance mechanism (*data not shown*). This resulted in hepatic OVA expression that was much lower than that in *Alb^sOVA^* mice (Fig. S1K). This level triggered robust OT-II proliferation in liver-draining LNs with no evidence of T cell deletion (Fig. 1H, I). These data together indicate that the liver, a tissue with tolerogenic nature^10–12^, has a unique mechanism that distinguishes high-dose from low-dose self-antigen and selectively deletes T cells recognizing high-dose antigen at homeostasis. Notably, this liver-specific tolerance mechanism was disrupted when the liver was perturbed by the hepatotoxic agent carbon tetrachloride (Fig. 1J).

Given that pTreg cell induction is the dominant peripheral tolerance program for intestinal self-antigens^7^, we next examined whether hepatic and pancreatic self-antigens utilize a similar program by assessing OT-II cell fate in *Alb^tmOVA^, Ptf1a^sOVA^, Ptf1a^tmOVA^* and *Ins1^sOVA^* mice in the shared LNs. The frequencies of T-BET^+^ Th1 cell were consistently highest in the liver LN (livLN), which receives the greatest lymphatic drainage from the liver and pancreas (Fig. S2A, B). FOXP3^+^ pTreg cell percentages also were highest in livLN when OVA originated from the liver, but also when it came from exocrine pancreas. This was in contrast to when OVA was expressed in pancreatic β cells, which led to the highest abundance of pTreg cells in duodenal LN (duoLN) (Fig. 1K), mirroring the distribution of pTreg cells specific for native islet antigens^6^. We hypothesized that this difference in response to exocrine versus endocrine antigen may be because if antigen is limiting, pTreg cell induction reflects antigen availability and that the use of STZ to release β-cell antigen overcame this limitation, revealing the superior pTreg induction potential of the duoLN. Indeed, when we used cerulein to acutely damage the exocrine pancreas in *Ptf1a^sOVA^* mice and this released more antigen, we observed a pTreg cell pattern similar to that seen with endocrine sOVA (Fig. 1L).

Overall, our data show that each tissue, depending on its physiological function and immune characteristics, employs distinct tolerance mechanisms toward self-antigens from different subcellular compartments. These factors, together with the amount of antigen delivered to LNs, dictate T cell fates at homeostasis.

### Type 1 and but not type 2 intestinal infections shifts exocrine-pancreas-reactive T cell fate in LNs shared between the pancreas and duodenum

Intestinal infections have been shown to reprogram T cell responses against dietary and self-antigens in the gut^7,13^. We previously demonstrated that certain intestinal viral infections, such as with reovirus strain type 1 Lang (T1L) and murine norovirus, can render islet-specific T cells more pro-inflammatory without spreading beyond the intestine^6^. This occurs via changes in the transcriptional profile of cDC1s through LN co-drainage^6^.. cDC1s are the main migratory dendritic cell population in the pancreas and liver and sample islet antigens^6^, and are also the primary DC subset that induces Th1 and CTLs. By contrast, migratory cDC2s, abundant in the gut, promote Th2 and Th17 differentiation but are virtually absent in pancreas (and liver). Based on this, we hypothesized that only cDC1-dependent, type 1 intestinal infections would alter T cell outcomes in the pancreas. To test this, we first infected *Ins1^sOVA^* and *Ptf1a^sOVA^*mice with the helminth *S. venezuelensis*, a type 2 infection. In both models, *S. venezuelensis* had no impact on the frequencies of GATA3^+^ OT-II cells and did not reduce Treg cells (Fig. S2C-F), as opposed to the response when the self-antigen originates from the gut^7^. By contrast, when we inoculated *Ins1^sOVA^* perorally (a term used interchangeably here with intragastric infection using a gavage needle) with T1L, we observed a marked increase in OVA-specific Th1 cells and a corresponding decrease in pTreg cells in the celiac (cel-) and duo-LNs (Fig. S2G, H), consistent with our previous findings with endogenous islet antigens^6^.

To determine whether this LN-co-drainage-driven regulation of endocrine pancreas-reactive T cells can be extended to the exocrine pancreas, we inoculated *Ptf1a^sOVA^* and *Ptf1a^tmOVA^*mice perorally with T1L. We again observed increased Th1 cell differentiation of OT-II cells in the same co-drained LNs, although with variable concomitant reduction in pTreg cell numbers (Fig. 2A-C, Fig. S3A-C). In parallel, OT-I cells in these LNs of T1L-infected mice acquired enhanced cytotoxic features, including expression of granzyme B (GzmB), interferon-γ (IFNγ), and tumor necrosis factor-α (TNFα) (Fig. 2D-G, Fig. S2A, S3D-G). We also used OVA tetramers to identify endogenous OVA-specific T cells. Again, we observed an increase in GzmB^+^CD8^+^ and T-BET^+^CD4^+^ T cells and a modest reduction in FOXP3^+^CD4^+^ T cells among tetramer^+^ cells in the co-drained LNs (Fig. 2H, I, Fig. S3H-J). Notably, in both the OT-I/OT-II and endogenous OVA-specific T cell contexts, this pro-inflammatory shift was absent in the livLN (Fig. 2A, D, Fig. S3H), a major pancreatic LN that however does not drain the gut. This finding highlights the specificity of the inter-organ crosstalk in that it occurs only in LNs that are co-drained by the tissue expressing the antigen and the tissue being perturbed. Furthermore, we hypothesized that LN co-drainage would only influence T cell fate for antigens that are actively being surveyed. Consistent with this, OT-II cells remained non-proliferative even in T1L-infected *Ptf1a^cOVA^* mice (Fig. S3K). These findings indicate that when an exocrine pancreatic self-antigen is available for immune presentation and recognition, intestinal infection can modulate exocrine-pancreas-specific T cell immunity, promoting inflammatory phenotypes.

**Figure 2.**
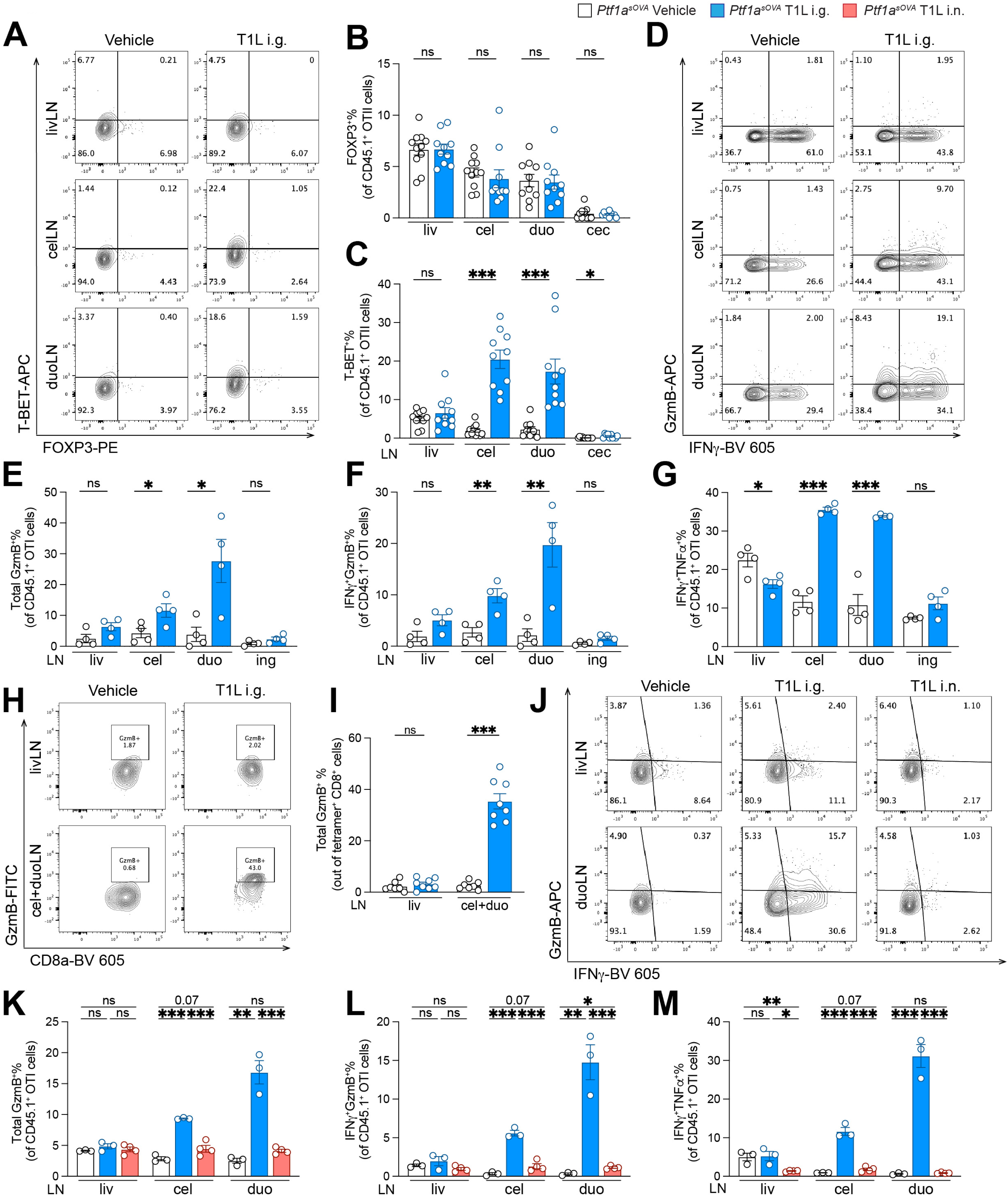
Type 1 and but not type 2 intestinal infections shifts exocrine-pancreas-reactive T cell fate by LN co-drainage. **A.** Flow plots of OT-II cells in LNs of *Ptf1a^sOVA^* mice inoculated perorally with T1L or vehicle control. **B-C.** Frequencies of FOXP3^+^ (**B**) or T-BET^+^ (**C**) among total OT-II cells in indicated LNs 72 h after transfer into mice in (**A**) (*n* = 10-11 per group, as indicated by symbols). **D.** Flow plots of OT-I cells in LNs of *Ptf1a^sOVA^* mice inoculated perorally with T1L or vehicle control. **E-G.** Frequencies of Total GzmB^+^ (**E**), IFNγ^+^GzmB^+^ (**F**) or IFNγ^+^TNFα^+^ (**G**) among total OT-I cells in indicated LNs 72 h after transfer into mice in (**D**) (*n* = 4 per group). **H-I**. Flow plots (**H**) and frequencies of GzmB^+^ among total OVA-tetramer^+^CD8^+^ T cells (**I**) in LNs of T1L-infected or vehicle-infected *Ptf1a^sOVA^* mice (*n* = 7-8 per group, as indicated by symbols). **J.** Flow plots of OT-I cells in LNs of *Ptf1a^sOVA^* mice that were either intragastrically or intranasally infected with T1L. **K-M.** Frequencies of Total GzmB^+^ (**K**), IFNγ^+^GzmB^+^ (**L**) or IFNγ^+^TNFα^+^ (**M**) among total OT-I cells in the indicated LNs 72 h after transfer into mice in (**J**) (*n* = 3-4 per group, as indicated by symbols). Data are representative of two independent experiments in B, C, E-G. Data represent one experiment in J, K-M. * p<0.05, ** p<0.01, *** p<0.001 by t-test.

To further solidify the requirements for LN co-drainage and regionality of the infection, we inoculated *Ptf1a^sOVA^* mice intranasally with T1L^14^, a route that targets the lung, an organ not co-drained with the pancreas. As anticipated, we observed increased CTLs cells among endogenous CD8⁺ T cells in the mediastinal LN (mtLN), confirming successful infection (Fig. S3L). However, we did not detect changes in the frequencies of CTLs in the pancreatic LNs of intranasally inoculated mice, in contrast to the robust response observed following peroral inoculation (Fig. 2J-M). Similarly, OVA-specific Th1 cells did not increase in the pancreatic LNs (Fig. S3M, N), despite intranasal T1L inoculation inducing more Th1 cells in the polyclonal compartment in the mtLN (Fig. S3O).

Together, these findings establish the requirements for immune crosstalk between gut and pancreas through LN co-drainage^6^, while expanding it to the exocrine pancreas.

### Intestinal reovirus-induced exocrine pancreas-reactive pro-inflammatory T cells migrate to the pancreas, leading to exocrine tissue destruction

To determine whether the T cells activated by reovirus T1L in LNs migrate to the pancreas and mediate pathological effects, we inoculated *Ptf1a^sOVA^* mice perorally with T1L at the time of OT-I and OT-II co-transfer. Immune cells isolated from the pancreas 7 days later had a substantial increase in the percentage and total number of OT-I and OT-II cells (Fig. 3A, B, S4A, B). In parallel, we detected increased infiltration of endogenous CD4⁺ and CD8⁺ T cells (Fig. S4C, D). Phenotypic analysis showed a higher proportion of GzmB^+^ OT-I cells and T-BET^+^ OT-II cells (Fig. 3C, D), indicating an inflammatory signature in the pancreas. Similar trends were observed among endogenous T cells, with a significantly higher percentage of CD8⁺ T cell expressing GzmB and an elevated Th1/pTreg cell ratio (Fig. S4E-I). Immunofluorescence staining confirmed widespread infiltration of both OVA-specific and polyclonal T cells into the pancreatic parenchyma (Fig. 3E). Histological analysis showed severe tissue damage, characterized by immune cell infiltration, necrosis, and interstitial edema (Fig. 3F, G). The percentage of apoptotic cells in the pancreas following T1L infection also was significantly increased (Fig. 3H, I). Of note, in the gut T1L infection in combination with OVA reactive cell transfer is not sufficient to cause measurable pathology^7^, suggesting that the pancreas, perhaps because it is inherently less regenerative than the gut epithelium, is more susceptible to damage. However, these pancreatic effects were not observed in the absence of OVA-specific T cell transfer (Fig. 3F, G), suggesting that while exogenous OT-I and OT-II transfer may overwhelm any pre-existing OVA-specific Treg cells in the pancreas, it remains a strong peripheral tolerance program to prevent pancreatic auto-destruction in responses to gastrointestinal infections.

**Figure 3.**
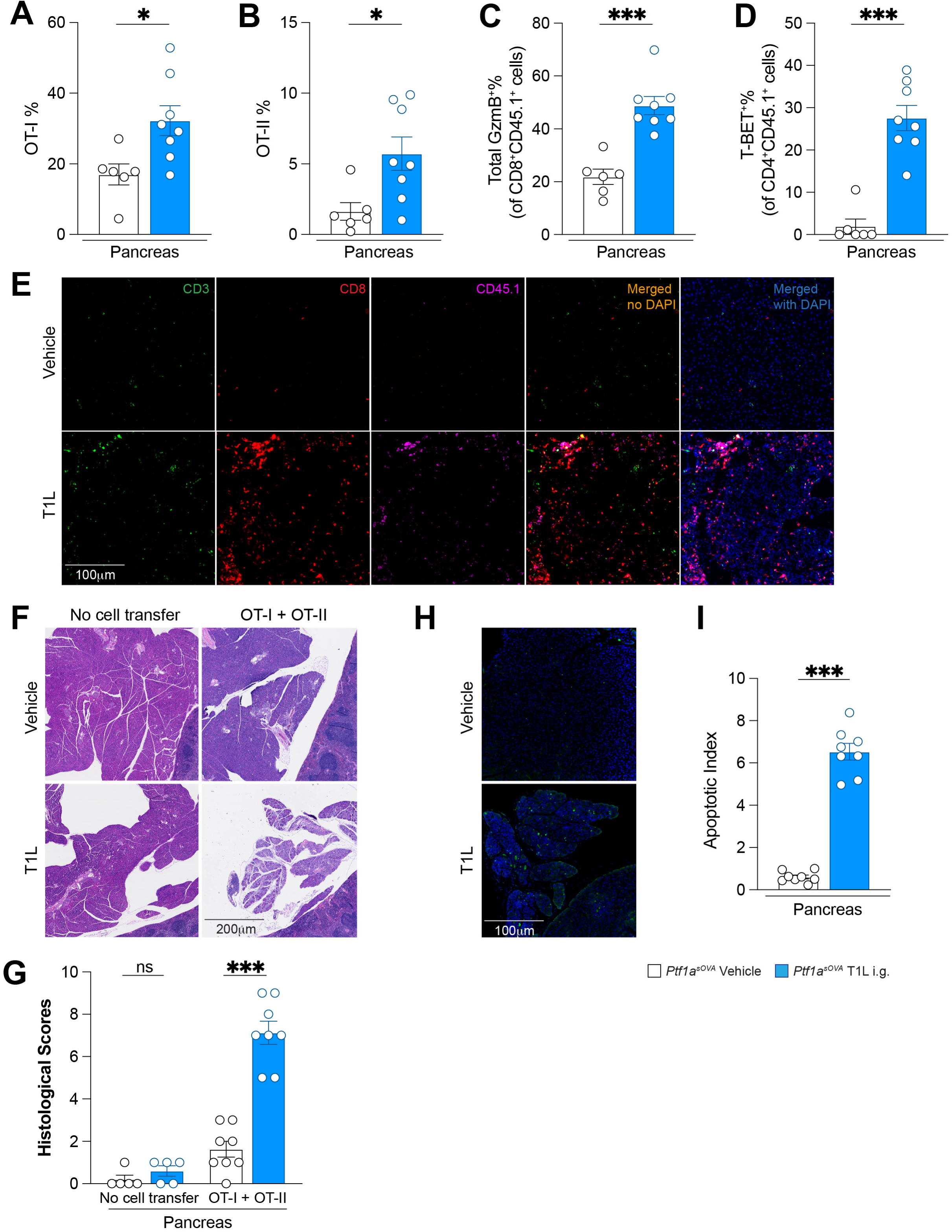
Intestinal reovirus-induced pro-inflammatory T cells infiltrate the pancreas and cause tissue destruction. **A-D.** Frequencies of OT-I cells among total CD8^+^ cells (**A**), OT-II cells among total CD4^+^ cells (**B**), GzmB^+^ among total OT-I cells (**C**) and T-BET^+^ among total OT-II cells (**D**) in the pancreas of orally T1L-infected or vehicle-infected *Ptf1a^sOVA^*mice 7 days post transfer (*n* = 6-8 per group, as indicated by symbols). **E.** Frozen sections of the pancreas from T T1L-infected or vehicle-infected *Ptf1a^sOVA^* mice 14 days post transfer stained for CD3, CD8, CD45.1 and DAPI (nuclei). Scale bar represents 100 μm. **F-I.** Sections of the pancreas from T1L-or mock-infected *Ptf1a^sOVA^*mice 14 days post transfer stained with H&E (**F**) or stained for apoptotic cells and DAPI (**H**). Scale bars represent 100 μm and 200 μm, respectively. Quantification of histological score (**G**) and apoptotic index (**I**) (*n* = 5-8 per group, as indicated by symbols). Data are pooled from two independent experiments in A-D, G, I. * p<0.05, ** p<0.01, *** p<0.001 by t-test.

These findings demonstrate that the intestinal reovirus infection-induced increase of inflammatory pancreas-reactive T cells in co-drained LNs correlates with the subsequent emergence of these activated autoreactive T cells in the pancreas where they mediate tissue destruction in the absence of pancreatic infection^6^.

### Intestinal reovirus leads to accelerated tumor progression in a genetic PDAC model

PDAC, the most common cancer of the exocrine pancreas, is an aggressive malignancy characterized by poor prognosis and low lymphocyte infiltration^8^. The PDAC microenvironment is immunosuppressive, with increased tumor-associated macrophages and Treg cells and diminished Th1 and CTL cell numbers, contributing to impaired anti-tumor immunity^15^. We therefore hypothesized that LN co-drainage could be exploited to modulate disease outcome by rewiring T cells recognizing tumor cells. To this end, we interbred KPC mice (*Ptf1a^Cre-ERT2/w^p53^fl/fl^Kras^G12D^*) with sOVA mice to produce animals in which sOVA is expressed in transformed pancreatic acinar cells (PDAC-sOVA mice). Tumor initiation was induced with tamoxifen containing diet at 3.5 weeks of age, followed by a 14-day period of tumor establishment. At that point, we adoptively transferred OVA-specific T cells into the mice, inoculated them perorally with reovirus T1L and either assessed T cell fate in the pancreatic LNs 3 days later or tumor burden 3 weeks later (Fig. 4A). As observed in *Ptf1a^sOVA^* mice, T1L robustly induced the generation of cytotoxic OT-I cells (Fig. 4B) and T-BET^+^ OT-II cells at the expense of pTreg cells in the co-drained LNs of PDAC-sOVA mice (Fig. 4C, D), showing that intestinal infection can induce type 1 immunity to pancreatic neoantigen in the context of PDAC. Perhaps unexpectedly, however, T1L infection was associated with significantly accelerated tumor progression, as assessed by the percentage of tumor area within the pancreas (Fig. 4E-G; Fig. S4A). This effect was also evident without transferring OVA-recognizing T cells, suggesting that the endogenous repertoire of OVA-reactive cells was sufficient to mediate accelerated tumor progression in the presence of LN perturbation. T1L-infected mice that received OT-I or both OT-I and OT-II cells showed the most severe pathology, with 95-100% lesion area (Fig. 4F, G), indicating that CD8⁺ T cells are primary mediators of this effect. To directly test the impact of CD8⁺ T cells on the oncogenesis of PDAC-sOVA mice, we administered a CD8β-neutralizing antibody beginning 7 days after T1L inoculation, to allow time for viral clearance, and continued antibody administration every 4 days. CD8-depleted mice showed significantly reduced tumor burden relative to isotype-treated controls (Fig. 4H, I), confirming that CD8⁺ T cells are the primary mediators of reovirus-mediated tumor acceleration. We concluded that the tumor-promoting effect of T1L in the genetic PDAC model was caused by the ubiquitous expression of OVA, which rendered nearly all pancreatic acinar cells susceptible to attack by OVA-specific T cells, including those that had not yet undergone full transformation and pushing them toward dedifferentiation. In addition, fully transformed pancreatic carcinoma cells employ multiple immunosuppressive mechanisms, making them less susceptible to T cell-mediated killing than pre-neoplastic or early-stage cells. However, these results underscore the capacity of intestinal viral infection to reshape pancreatic T cell immunity through shared LNs, with a potent influence on PDAC progression.

**Figure 4.**
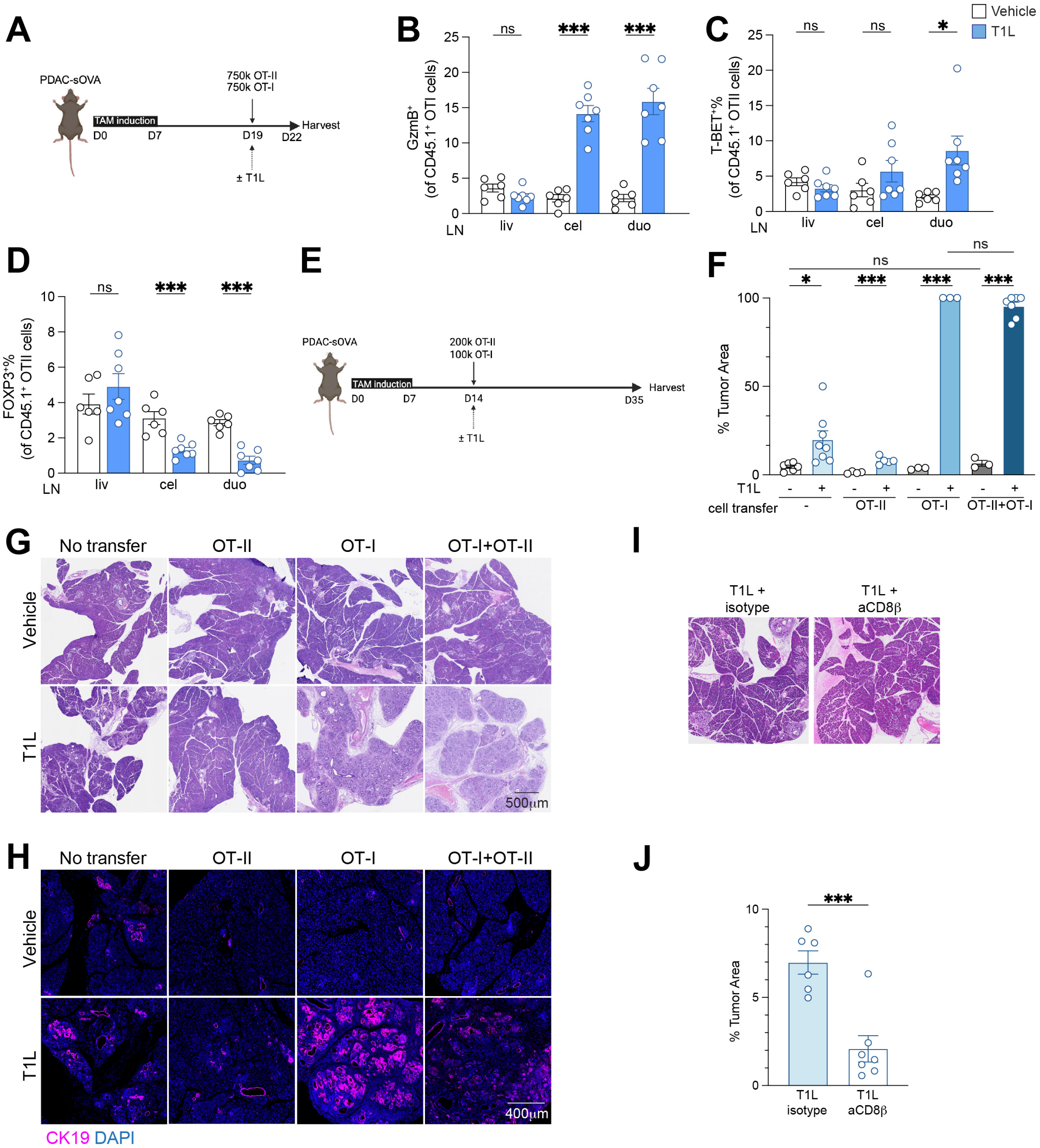
Intestinal reovirus infection exacerbates pancreatic tumor growth in a genetic PDAC model by a CD8^+^ T cell-dependent manner. **A**. Experimental scheme for tumor development, cell transfer and detection of T cell fate in the LNs. **B-D**. Frequencies of GzmB^+^ among total OT-I cells (**B**), T-BET^+^ (**C**) or FOXP3^+^ (**D**) among total OT-II cells in indicated LNs 72 h after co-transfer into T1L-infected or vehicle-infected PDAC-sOVA mice (*n* = 6-7 per group, as indicated by symbols). **E.** Experimental scheme for tumor development, cell transfer and analysis of tumor area. **F-H**. Sections of pancreas from PDAC-sOVA mice with indicated cell transfer and T1L infection status stained with H&E (**G**) or CK19 antibody (**H**). The percentage of tumor area from each section was quantified (**F**) (*n* = 3-8 per group, as indicated by symbols). Scale bar represents 500 μm (**G**) and 400 μm (**H**). **I-J**. Sections of pancreas from T1L-infected PDAC-sOVA mice treated with a CD8β neutralizing antibody or isotype control stained with H&E (**I**). The percentage of tumor area from each section was quantified (**J**) (*n* = 6-7 per group, as indicated by symbols). Scale bar represents 400 μm. Data are pooled from two independent experiments in B-D, F, J. * p<0.05, ** p<0.01, *** p<0.001 by t-test.

### Intestinal reovirus infection enhances T cell infiltration and tumor control in an orthotopic PDAC model

In human PDAC, tumor initiation occurs neither uniformly in all acinar cells simultaneously, nor do all acinar cells express neoantigens. Therefore, to model a scenario in which only tumor cells express a neoantigen and not all acinar cells have transformation potential, we employed an orthotopic system using a syngeneic KPC-derived tumor cell line^16^. This system also permitted us to easily test if the subcellular compartment in which a tumor antigen is expressed dictates the capacity of the adaptive immune system to exert anti-tumor control. To this end, we stably transduced KPC cells with either a control lentiviral vector (empty vector, EV) or a vector encoding sOVA, cOVA or tmOVA, whereby sOVA and cOVA were expressed at similar protein levels and therefore used for subsequent analyses (Fig. S5A). The KPC cells were injected into the pancreas of B6 mice, and tumors were allowed to develop for 11 days. In the absence of further experimental interference, all three KPC lines gave rise to similarly sized tumors (Fig. S5B, C). To confirm antigen drainage and T cell priming in the LNs, mice were administered OT-I and OT-II cells and inoculated perorally with reovirus T1L or not (Fig. 5A). We observed robust proliferation of OT-II cells in pancreatic LNs only when KPC-sOVA but not when KPC-cOVA cells were injected, with OT-II cells in livLN showing the highest proliferation, followed by celLN and duoLN (Fig. S5D), mirroring the previously mapped pancreatic drainage^6^. Although proliferation did not significantly change following T1L infection (Fig. S5D), we detected increased frequencies of GzmB⁺ OT-I and T-BET⁺ OT-II cells in co-drained LNs, indicating that T1L infection led to inflammatory T cell polarization in the orthotopic setting (Fig. 5B, C, S5E-H).

**Figure 5.**
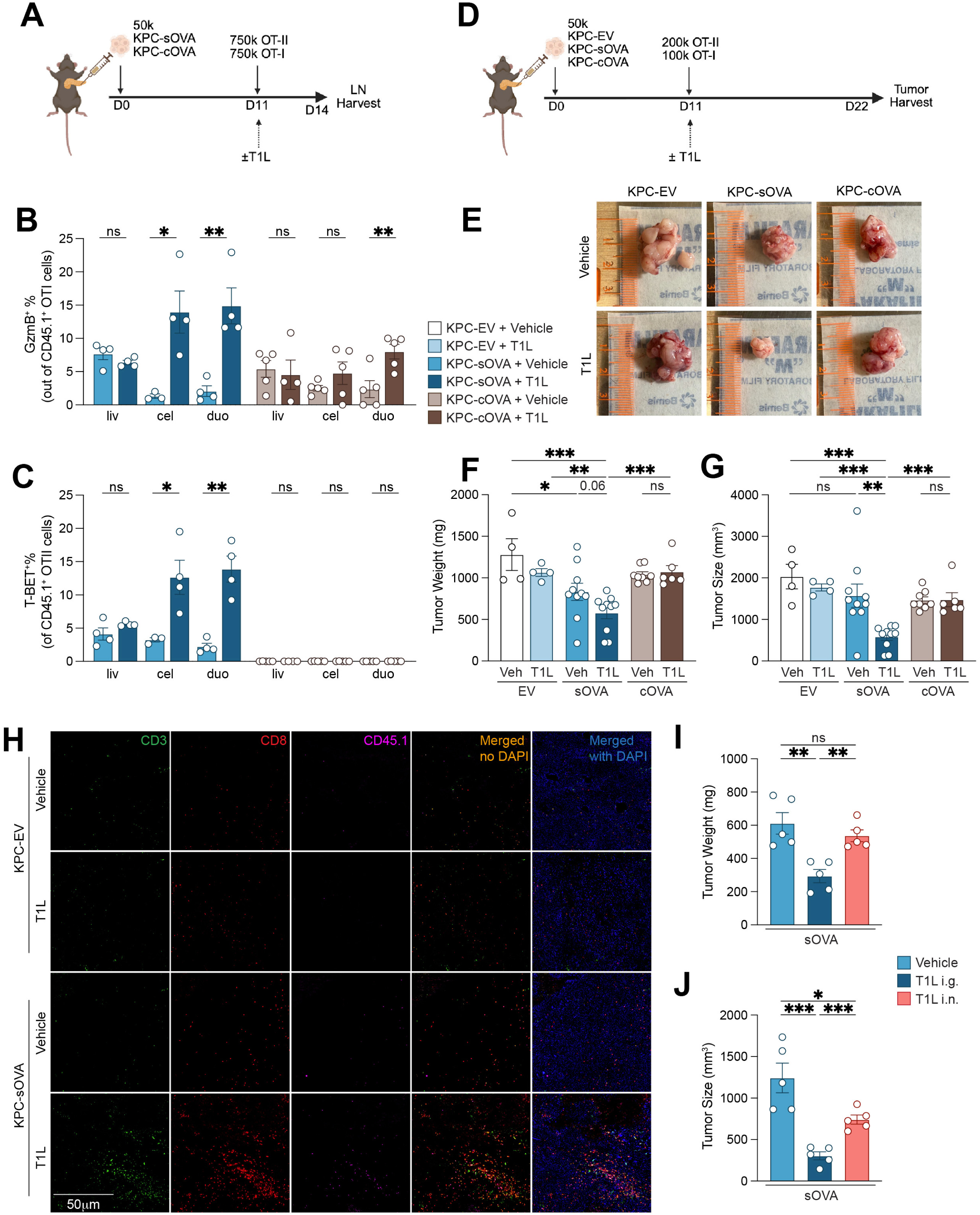
Intestinal reovirus infection enhances T cell infiltration and tumor control in an orthotopic PDAC model. **A, D**. Schematic of orthotopic PDAC model for assessing T cell fate in the LNs (**A**) and tumor progression (**D**). **B-C**. Frequencies of GzmB^+^ among total OT-I cells (**B**) or T-BET^+^ among total OT-II (**C**) cells in indicated LNs of mice receiving sOVA- or cOVA-expressing tumor cells (*n* = 4-5 per group, as indicated by symbols). **E.** Representative photographs of tumors removed from mice with indicated tumor implants and T1L infection status. **F-G.** Weights (**F**) and sizes (**F**) of tumors from mice in (**E**) (*n* = 4-10 per group, as indicated by symbols). **H.** Tumor frozen sections from mice in (**E**) were stained for CD3, CD8, CD45.1 and DAPI (nuclei). Scale bar represents 50 μm. **I-J.** Weights (**I**) and sizes (**J**) of tumors from mice receiving sOVA-expressing KPC cells intragastrically or intranasally infected with T1L (*n* = 5 per group). Data are pooled from two independent experiments in F, G. Data represent one experiment in B, C, I, J. * p<0.05, ** p<0.01, *** p<0.001 by t-test.

We next tested the effect of T1L infection on tumor control. Mice were injected orthotopically with either 50,000 KPC-EV, KPC-sOVA or KPC-cOVA cells. On day 11, when tumors were similarly established in all groups (Fig. S5B, C), mice were administered OT-I and OT-II cells and inoculated perorally with T1L (Fig. 5D). T1L infection reduced tumor burden in KPC-sOVA but not KPC-cOVA or KPC-EV tumor-bearing mice relative to the uninfected controls (Fig. 5E-G), in congruence with the observations in the LNs where KPC-cOVA elicited no OVA-specific T cell responses (Fig. 5B, C). Immunofluorescence staining revealed significantly increased infiltration of both OVA-specific and endogenous T cells in the pancreas of T1L-infected mice with KPC-sOVA tumors but not KPC-EV or KPC-cOVA mice (Fig. 5H and *data not shown*). To confirm that LN co-drainage was required for this effect, we compared tumor outcomes in mice inoculated with T1L either intragastrically or intranasally. Intragastric inoculated showed superior tumor control than that following intranasal inoculation, supporting the requirement for direct gut-to-pancreas immune interaction for this modulation (Fig. 5I, J).

In a final series of experiments, we determined whether peroral administration of a tumor-associated neoantigen could enhance anti-tumor immunity in the presence of an intestinal infection (Fig. S5I). A single dose of oral OVA was sufficient to increase the number of GzmB⁺ OVA-specific T cells in pancreatic draining LNs (Fig. S5J, K), which reduced tumor size, particularly in T1L-infected mice with KPC-sOVA tumors (Fig. S5L, M). Given the aggressive nature of this tumor cell line, we hypothesized that the intervention might be more effective if the initial tumor burden was lower. Therefore, we repeated the experiment using a lower tumor inoculum (5,000 cells; Fig. S5I). In this setting, we again observed a reduction in tumor burden, with the T1L-infected, sOVA-tumor-bearing group having the most significant response, as approximately 25% of the mice achieved complete tumor clearance at the end point of the experiment (Fig. S5N, O). Together, these data demonstrate that intestinal infection can enhance tumor-specific T cell responses and reduce tumor burden in PDAC through LN co-drainage when the antigen is confined to the tumor cells.

### Intestinal reovirus infection renders liver-specific T cells more pro-inflammatory leading to liver damage

In our previous experiments we largely ignored the role of liver in tri-organ immune crosstalk, despite it being a major lymph-producing organ. To assess whether immune-modulation by intestinal type 1 infections extends to the liver, we perorally inoculated *Alb^sOVA^* and *Alb^tmOVA^* mice with T1L, followed by OT-II transfer. In *Alb^sOVA^* mice, T1L did not reverse OT-II clonal deletion (Fig. S6A, B), reinforcing our suspicion that this is a liver-intrinsic phenomenon. In *Alb^tmOVA^* mice however, we observed increased Th1 and reduced pTreg cell frequencies specifically in the celLN, the primary LN co-draining the liver and gut (Fig. 6A-C). In the CD8⁺ T cell compartment, OT-I cells from infected mice displayed an enhanced cytotoxic profile in both celLN and duoLN (Fig. 6D-G). We suspect that the duoLN response reflects a combination of greater CD8⁺ T cell sensitivity to antigen and possible shifts in hepatic drainage patterns under inflammatory conditions.

**Figure 6.**
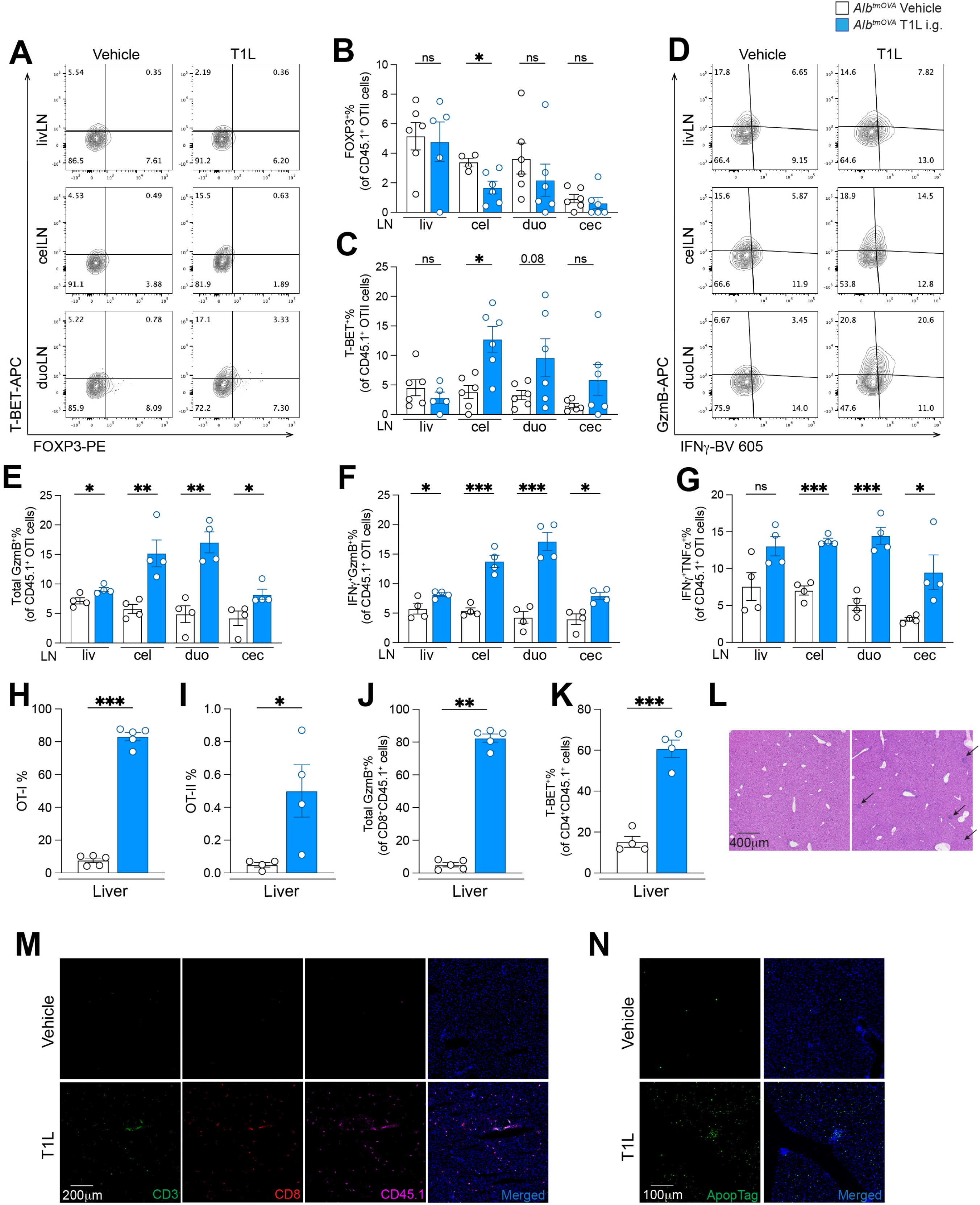
Intestinal reovirus infection induces liver-reactive T cells and hepatic damage by LN co-drainage. **A.** Plots of OT-II cells in indicated LNs of orally T1L-infected or vehicle-infected *Alb^tmOVA^* mice, representative of gating applies in (**B**) and (**C**). **B-C.** Frequencies of FOXP3^+^ (**B**) or T-BET^+^ (**C**) among total OT-II cells in indicated LNs 72 h after transfer into mice in (**A**) (*n* = 5-6 per group, as indicated by symbols). **D.** Plots of OT-I cells in indicated LNs of T1L-or vehicle-infected *Alb^tmOVA^* mice, representative of gating applied in (**E**) – (**G**). **E-G.** Frequencies of total GzmB^+^ (**E**), IFNγ^+^GzmB^+^ (**F**) or IFNγ^+^TNFα^+^ (**G**) among total OT-I cells in indicated LNs 72 h after transferred into mice in (**D**) (*n* = 4 per group). **H-K**. Frequencies of OT-I cells among total CD8^+^ cells (**H**), OT-II cells among total CD4^+^ cells (**I**), GzmB^+^ among total OT-I cells (**J**) and T-BET^+^ among total OT-II cells (**K**) in the liver of T1L- or mock-infected *Alb^tmOVA^* mice 7 days post transfer (*n* = 5 per group). **L.** Sections of the liver from orally T1L-infected or vehicle-infected *Alb^tmOVA^* mice 14 days post transfer stain with H&E. Scale bar represents 400 μm. **M.** Frozen sections of the liver from T1L-infected or vehicle-infected *Ptf1a^sOVA^* mice stained for CD3, CD8, CD45.1 and DAPI (nuclei). Scale bar represents 200 μm. **N.** Sections of the liver from T1L-infected or vehicle-infected *Alb^tmOVA^* mice 14 days post infection stained for apoptotic cells. Scale bar represents 100 μm. Data are representative of two independent experiments in B, C, E-K. * p<0.05, ** p<0.01, *** p<0.001 by t-test.

As seen in the pancreas, activated T cells also subsequently migrated to the liver. Both OT-I and OT-II cells accumulated in the liver following perioral T1L inoculation, with increases observed in both the percentage and absolute number of each (Fig. 6H, I, S6C, D). Endogenous CD8⁺ but not CD4⁺ T cells also infiltrated the liver in greater numbers (Fig. S6E, F). Phenotypic analysis using flow cytometry showed higher frequencies of hepatic GzmB⁺ OT-I and T-BET^+^ OT-II cells upon T1L infection (Fig. 6J-K). Endogenous CD8⁺ T cells also increased in number and cytotoxicity (Fig. S6E, G, H), while the effect on endogenous CD4⁺ T cells was more limited (Fig. S6F, I-K). Histological analysis of liver sections revealed localized immune cell aggregates accompanied by tissue damage (Fig. 6L). Immunofluorescence staining showed extensive infiltration of both OVA-specific transferred T cells and endogenous T cells throughout the hepatic parenchyma (Fig. 6L), while ApopTag staining demonstrated a marked increase in apoptotic cells, notably in areas densely populated by immune infiltrates (Fig. 6M).

Collectively, these findings demonstrate that intestinal reovirus infection impact T cell immunity not only in the endocrine^6^ and exocrine pancreas but also in the liver, revealing a gut-to-liver immune axis through LN sharing.

### Hepatotoxic concanavalin A induces pancreas- and gut-reactive Th1 cells and reduces pTreg cells

We finally sought to determine whether perturbations of the liver induce immune shifts in the pancreas or intestine through shared LNs, thus, if immunological liver-to-gut or -pancreas axes exist. Since human hepatitis viruses do not infect mice^17,18^, current mouse models for hepatitis viruses rely on humanized systems^17,19^ or other hepatotropic viruses that also infect the pancreas^20–22^. Therefore, we instead used a non-viral model: Concanavalin A (conA), which induces autoimmune hepatitis and causes liver-specific damage by a Kupffer cell- and T cell-dependent type 1 immune response^23^. We first tested the effect of conA on T cell responses in *Alb^tmOVA^* mice. Following conA administration, we observed significant liver injury (Fig. 7A) and increased OT-II proliferation in liver-draining LNs (Fig. S7A). Notably, pTreg cell induction was suppressed in livLN and celLN, and Th1 cells increased in these nodes (Fig. 7B, C), establishing conA as an agent for probing liver-driven immune regulation by LN co-drainage.

**Figure 7.**
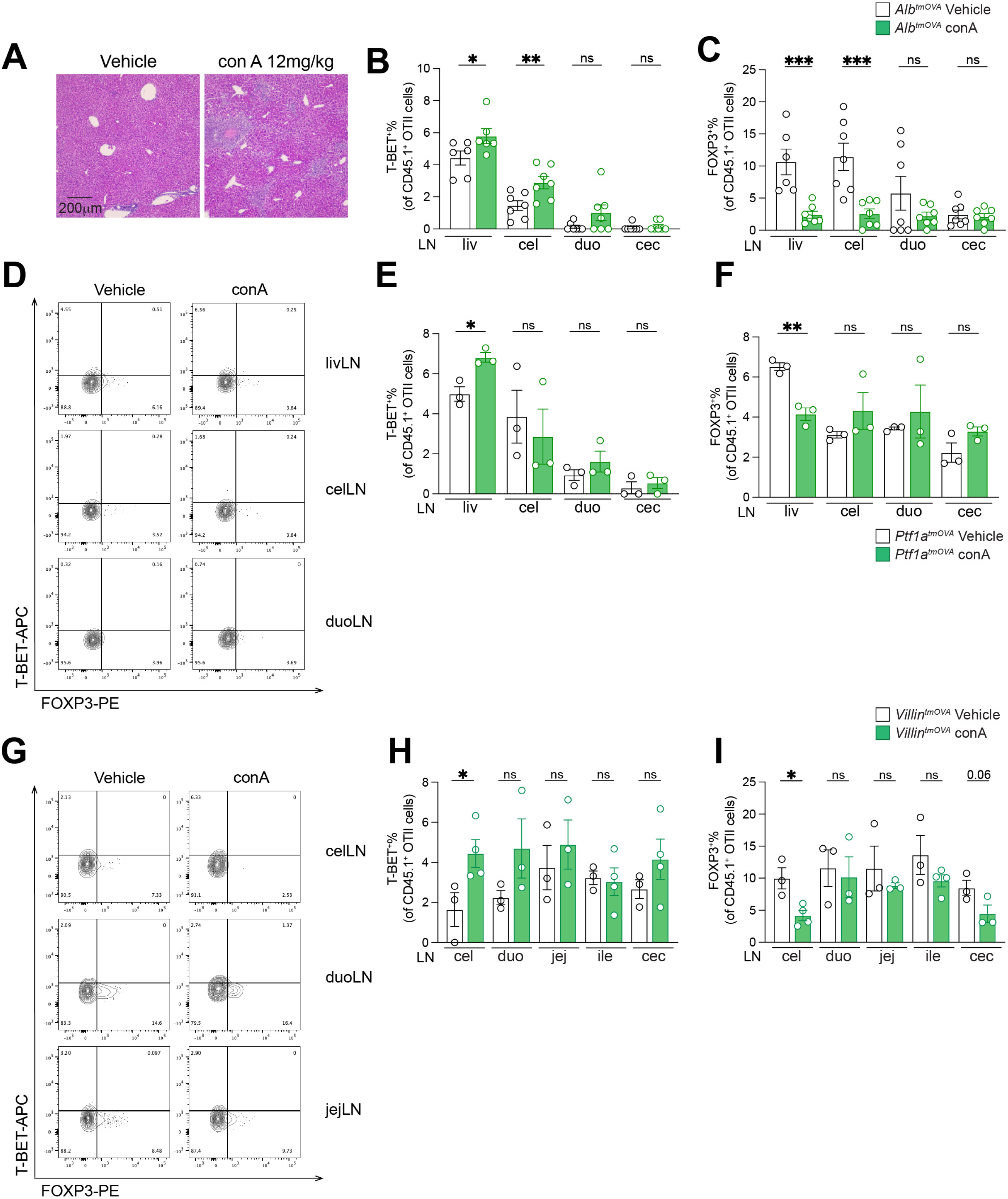
Con A-induced liver injury promotes pro-inflammatory conversion of pancreas-and gut-reactive T cells. **A.** Sections of the liver from con A- or vehicle-treated *Alb^tmOVA^* mice stained with H&E. Scale bar represents 200 μm. **B-C.** Frequencies of T-BET^+^ (**B**) or FOXP3^+^ (**C**) among total OT-II cells in indicated LNs 96 h after transfer into mice in (**A**) (*n* = 6 per group). **D.** Plots of OT-II cells in indicated LNs of con A- or vehicle-treated *Ptf1a^tmOVA^*mice, representative gating applied in (**E**) and (**F**) (*n* = 3 per group). **E-F.** Frequencies of T-BET^+^ (**E**) or FOXP3^+^ (**F**) among total OT-II cells in indicated LNs 96 h after transfer into mice in (**D**). **G.** Plots of OT-II cells in indicated LNs of con A- or vehicle-treated *Villin^tmOVA^* mice, representative gating applied in (**H**) and (**I**). **H-I.** Frequencies of T-BET^+^ (**E**) or FOXP3^+^ (**F**) among total OT-II cells in indicated LNs 96 h after transfer into mice in (**G**) (*n* = 3-4 per group, as indicated by symbols). Data are representative of two independent experiments in B, C, E, F, H, I. * p<0.05, ** p<0.01, *** p<0.001 by t-test.

To determine whether conA-induced liver inflammation alters T cell responses to pancreatic and intestinal self-antigens, we treated *Ptf1a^tmOVA^* or *Villin^tmOVA^* mice, which express the same antigen as *Alb^tmOVA^*, with conA and monitored T cell differentiation. In *Ptf1a^tmOVA^* mice, conA treatment led to an increase in Th1 cells and a reduction in pTreg cells within livLN (Fig. 7D-F), the LN with the highest shared drainage between liver and pancreas. Similarly, in *Villin^tmOVA^* mice, conA treatment led to an increase in Th1 cells and a reduction in pTreg cells, specifically within the celLN, which is the only LN shared by liver and duodenum (Fig. 7G-I). Comparable results were obtained in experiments using *Ptf1a^sOVA^* (Fig. S7B-C) and *Villin^sOVA^*(Fig. S7D-E) mice. Although pancreatic antigens are also drained into celLN and duoLN, and gut antigens are drained into multiple gut draining LNs, no significant T cell phenotypic changes were observed in those LNs. This result suggests that the observed effects are confined to the LNs co-drained with the liver and are not systemic, excluding the possibility that our observations are due to conA directly affecting T cells.

Thus, liver perturbation also reciprocally regulates T cell immunity of other co-drained tissues, establishing liver-to-gut/pancreas axes and highlighting LN co-drainage as a critical immunological mechanism for inter-organ immune crosstalk.

## Discussion

In this study, we uncovered that tissue-specific tolerance mechanisms and LN co-drainage collectively dictate T cell immune outcomes in the upper digestive system. By standardizing the antigen and T cell receptor affinity, and leveraging the fact that the liver, pancreas and duodenum drain into the same LNs, we disentangled the contribution of LN environment and tissue source to T cell polarization. Specifically, we discovered that liver and pancreas, which are functionally and immunologically different from the gut, adopt distinct tissue-specific mechanisms for maintaining tolerance to self-antigens from different subcellular compartments and with different dosages at homeostasis. Using an intestinal perturbation, we expanded on our previous findings^6^ and showed that one of tolerance programs, pTreg cell differentiation, can also be disrupted in the exocrine pancreas through LN sharing. While this disruption can result in pancreatic autoimmunity, it has the potential of being harnessed to enhance PDAC tumor control. Finally, our study identified reciprocal immune regulation between the liver and gut through LN co-drainage, establishing liver-pancreas-gut axes.

Maintaining tolerance to self-antigens is critical for preventing autoimmunity. Because central tolerance is incomplete^24^, multiple peripheral tolerance mechanisms have evolved to prevent aberrant responses to self. However, it remains unclear whether all tissues preferentially employ a common mechanism, or whether, analogous to inflammatory responses, they adapt their tolerance mechanisms to their physiological roles. We found that peripheral tolerance is sculpted jointly by tissue of origin and the subcellular localization of antigens. Cytosolic self-antigens in tissues with low cell turnover, like the pancreas and liver, are tolerated through immune ignorance, as antigen levels remained below the threshold required for T cell activation. In contrast, secreted or membrane-shed self-antigens preferentially induced pTreg cell differentiation, particularly in the gut^7^. Interestingly, high levels of secreted self-antigen from the liver triggered pruning of antigen-specific T cells, suggesting a liver-specific, dominant deletional strategy. These differences likely reflect tissue-specific immune pressures. The gut, continuously exposed to foreign antigens, relies heavily on active suppression to maintain immune quiescence, favoring pTreg cell induction, which enables flexible rewiring of immune responses during inflammatory events such as epithelial breach. The pancreas, physically connected to the duodenum and susceptible to pancreatic ductal reflux, may employ a similar mechanism to coordinate local immune responses with the gut. In contrast, the liver is largely sterile, so immune responses there may be primarily triggered by self-antigens, contributing to pathologies from unwanted pro-inflammatory reactions in autoimmune conditions^25,26^ or immunosuppressive outcomes such as chronic infections or tumor outgrowth^27^. Additionally, as a key site for producing highly abundant serum proteins and sampling blood-borne antigens of other origins^28^, clonal deletion may be the most efficient strategy for maintaining tolerance in the liver, although the precise mechanism or cellular contributors remain to be defined in future investigation. Thus, identical antigens can induce distinct T cell fates in the same LNs depending on tissue origins, underscoring that tissues employ specialized tolerance programs based on their antigen accessibility, local immune environment, and physiological function. Understanding these tissue-specific mechanisms has broad implications: Autoimmune diseases may emerge when these programs fail, while cancers may exploit them to evade immune detection. By defining how tolerance varies by tissue, we can design more precise immunotherapeutics that either reinforce these programs to prevent autoimmunity or temporarily override them to boost anti-tumor immunity.

Our study also expands the physiological relevance of LN co-drainage by establishing liver-pancreas-gut axes and demonstrating the reciprocal immune regulation between these organs. The bidirectional communication introduces an additional layer of immune modulation that must be considered alongside local and systemic factors. From a translational perspective, this mechanism presents exciting therapeutic opportunities. The gut, which is readily accessible, may serve as a non-invasive entry point for modulating immune responses in more secluded organs like the liver or pancreas through shared lymphatic drainage. This anatomic and immunologic connection opens the possibility of developing new strategies to either dampen unwanted autoimmune responses or enhance anti-tumor or anti-viral immunity by targeting the immune system at a regional level.

Another notable finding in this study is that the pancreas is markedly more susceptible to autoimmune destruction than the liver (Fig. 3 and Fig. 4) and gut^7^, even when exposed to the same number of antigen-specific T cells and equivalent inflammatory stimuli. Although differences in the number of infiltrating T cells per gram of tissue or variation in local tolerogenic milieu may contribute, these results also suggest that the consequences of LN co-drainage are not uniform across organs. Instead, they appear to be shaped by each tissue’s intrinsic regenerative and immunological capacities. The pancreas, with its limited regenerative ability compared to the liver or gut, appears particularly vulnerable to immune-mediated injury once an autoimmune response is initiated. Consequently, the inflammation mediated by shared LNs exerts more destructive and lasting effects on the pancreas than on more regenerative tissues. Physiologically, this distinction may help explain the higher incidence and chronicity of autoimmune diseases targeting the pancreas, such as type 1 diabetes, whereas comparable insults in the liver are more often transient or self-resolving. Exploring the impact of LN co-drainage on primary or metastatic liver tumors represents an interesting direction for future investigation.

In our experiments, genetically engineered and orthotopic PDAC mouse models had markedly different outcomes in response to elevated antigen-specific T cell infiltration (Fig. 4 and Fig. 5), highlighting the importance of choosing suitable models for studies of PDAC immunobiology. Genetically engineered mouse models (GEMMs), like the KPC model, recapitulate the gradual tumor evolution and desmoplastic stroma formation characteristic of human PDAC, but fail to capture the stochastic nature of tumor initiation in humans, in which only a subset of acinar cells gradually dedifferentiates and adopt ductal features. Since the pancreas is a tissue prone to stress^29^, acinar cells have the capacity to partially dedifferentiate under stress conditions^30,31^. In this context, indiscriminate killing by antigen-specific T cells, due to the ubiquitous expression of the model neoantigen, may exacerbate tissue stress and inadvertently accelerate acinar-to-ductal metaplasia, thereby worsening PDAC progression. In contrast, orthotopic transplantation mouse models offer synchronized tumor onset and tighter temporal control and limit the expression of neoantigens to tumor cells. As such, only malignant cells become targets of immune cells, allowing more specific T cell-mediated tumor cell killing. Our data also emphasize the importance of intervention timing. While boosting antigen-specific T cell infiltration can be beneficial, the therapeutic effect diminishes as the tumor burden increases. This phenomenon is likely due to physical and immunological barriers established by the stroma, including extracellular matrix deposition and exclusion of effector cells from tumor nests^32^. Moreover, advanced lesions may develop additional resistance mechanisms, such as antigen loss or immune checkpoint upregulation^32^. Collectively, our findings suggest that model selection and timing of immune intervention must be carefully aligned with the specific biological question being addressed in PDAC research.

Collectively, our study reveals the complexity of forces shaping adaptive immune outcomes in the upper digestive system, offering insights into hitherto idiopathic diseases like autoimmunity and cancer, as well as their heterogeneity and interindividual variability.

## Limitations of the study

Our studies rely on a model antigen, OVA. While we were able to confirm key results using tetramers that detect endogenous OVA-reactive T cells, it is likely that the mechanisms we reveal are not as active as when we deliberately transfer OVA reactive T cells. This is a general limitation in adoptive transfer systems used in immunology. We concentrated the current study to exploring the consequences of the combined effect of LN co-drainage and antigen source to pancreatic diseases, leaving understanding the liver’s potentially unique deletion-inducing mechanisms to handle high dose self-antigens and the implications of LN co-drainage for liver disease for future investigations.

## Acknowledgements

We thank the University of Chicago Animal Resource Center and microscopy, flow cytometry, functional genomics cores. We thank the members of the Esterhazy lab for invaluable discussions and suggestions along the study. This work was funded by the National Institutes of Health (NIH) R01 DK133393, NIH R01 AI038339, the Searle Scholar’s Program, Pew Charitable Trust, and University of Chicago start-up funds (to D. Esterházy). Additional support was provided by NIH T32 AI007090 (to M. R. Komnick), Alba Tull and the Tull Family Foundation (G. M. Taylor), and the Heinz Endowments (to T. S. Dermody).

## Author contributions

Conceptualization, Y.Z., H.B., and D.E.; Methodology, Y.Z., P.W., H.B., E.S., M.R.K. C.S., A.M., and D.E.; Investigation, Y.Z., P.W., H.B., E.S., M.R.K., G.T., K.F., S.L., C.S., A.M., and D.E.; Validation, Y.Z., P.W., H.B., E.S., M.R.K., and D.E.; Formal Analysis, Y.Z., P.W., E.S., M.R.K., and D.E.; Data Curation, Y.Z.; Writing – Original Draft, Y.Z.; Visualization, Y.Z.; Writing – Review and Editing, Y.Z., P.W., H.B., E.S., G.T., K.F., M.R.K., C.S., A.M., T.S.D., and D.E.; Funding Acquisition, T.S.D, A.M., and D.E.; Resources, T.S.D., A.M., and D.E.; Supervision, D.E.

## Materials and Methods

### KEY RESOURCES TABLE

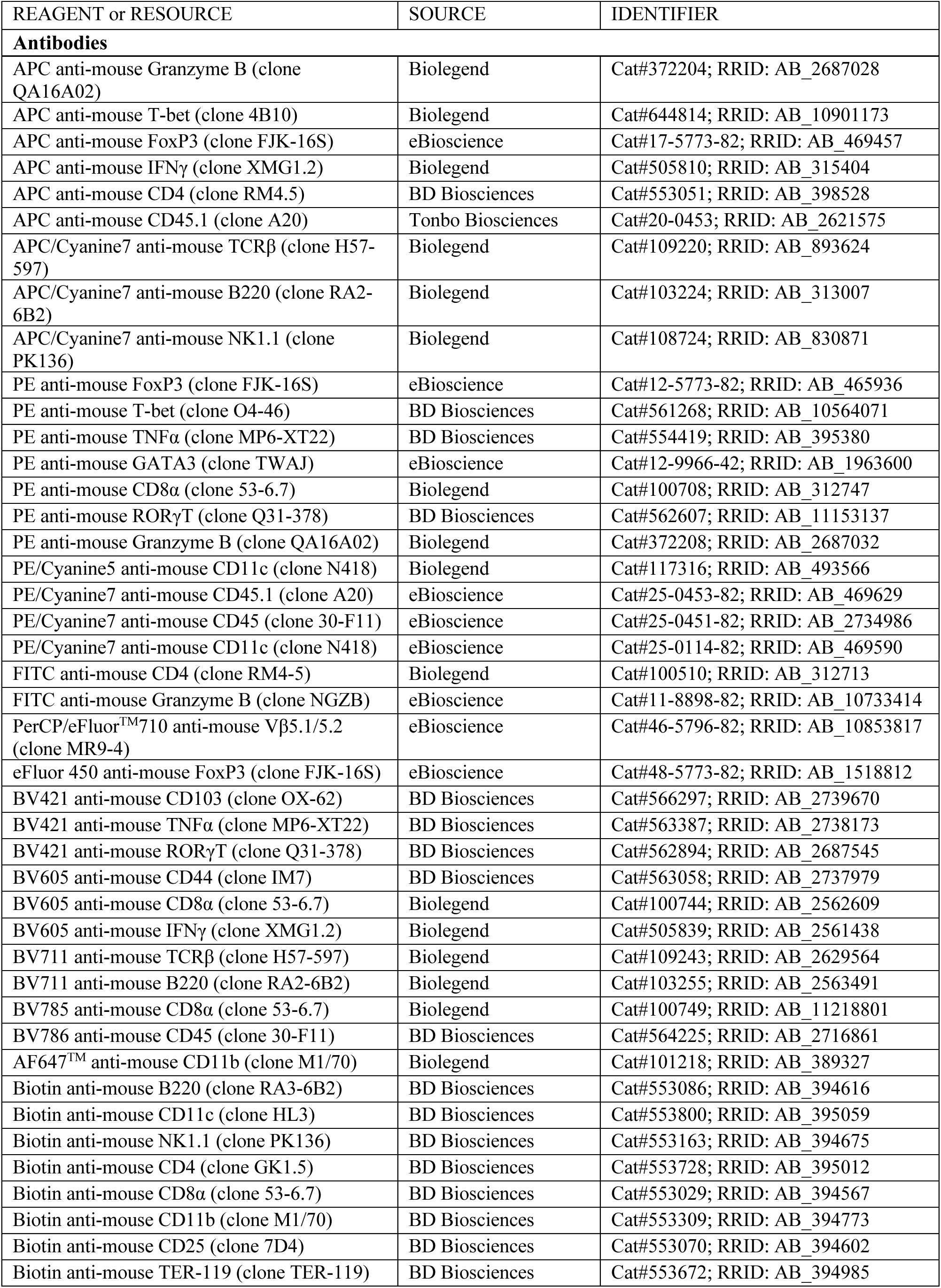
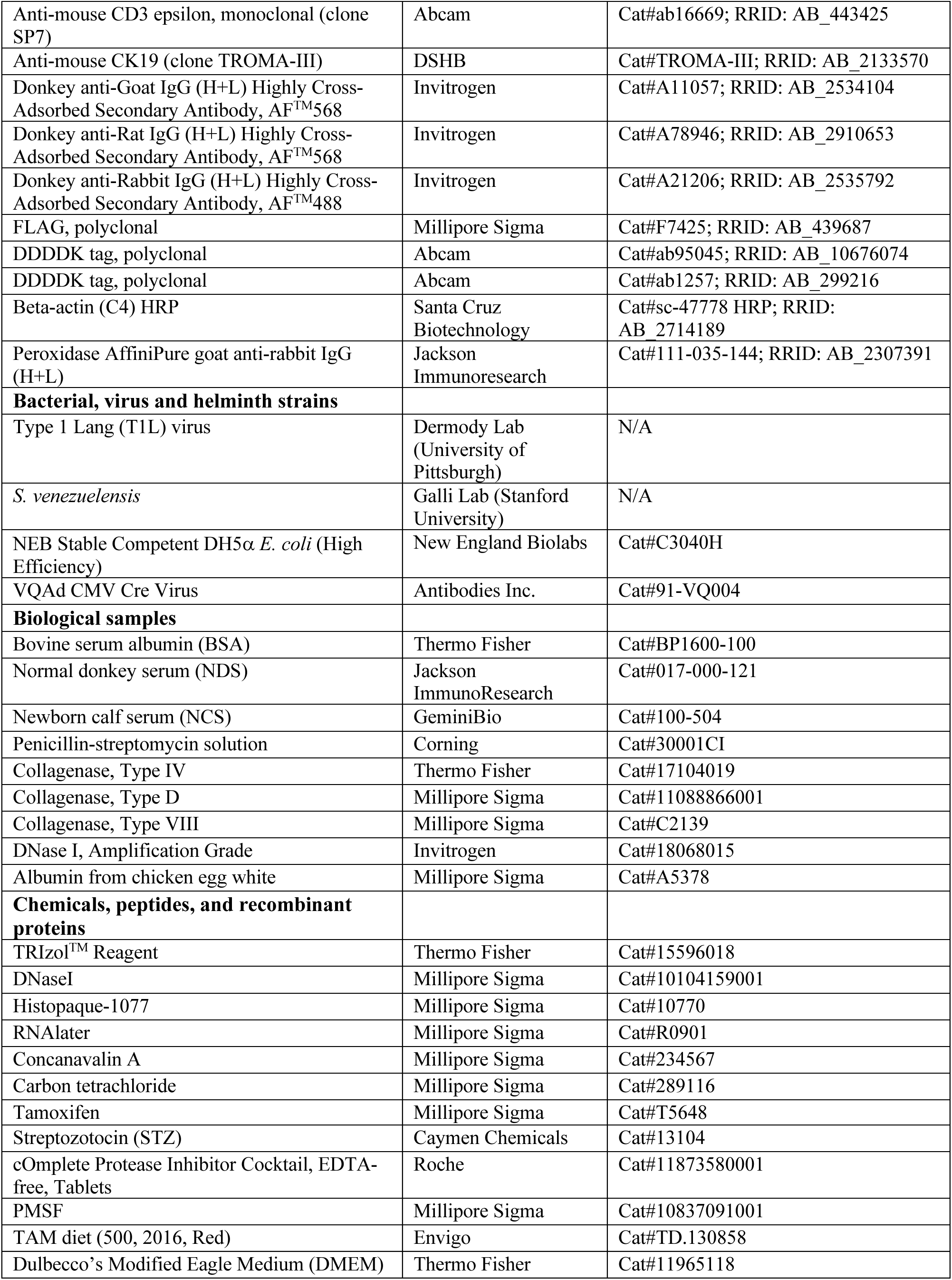
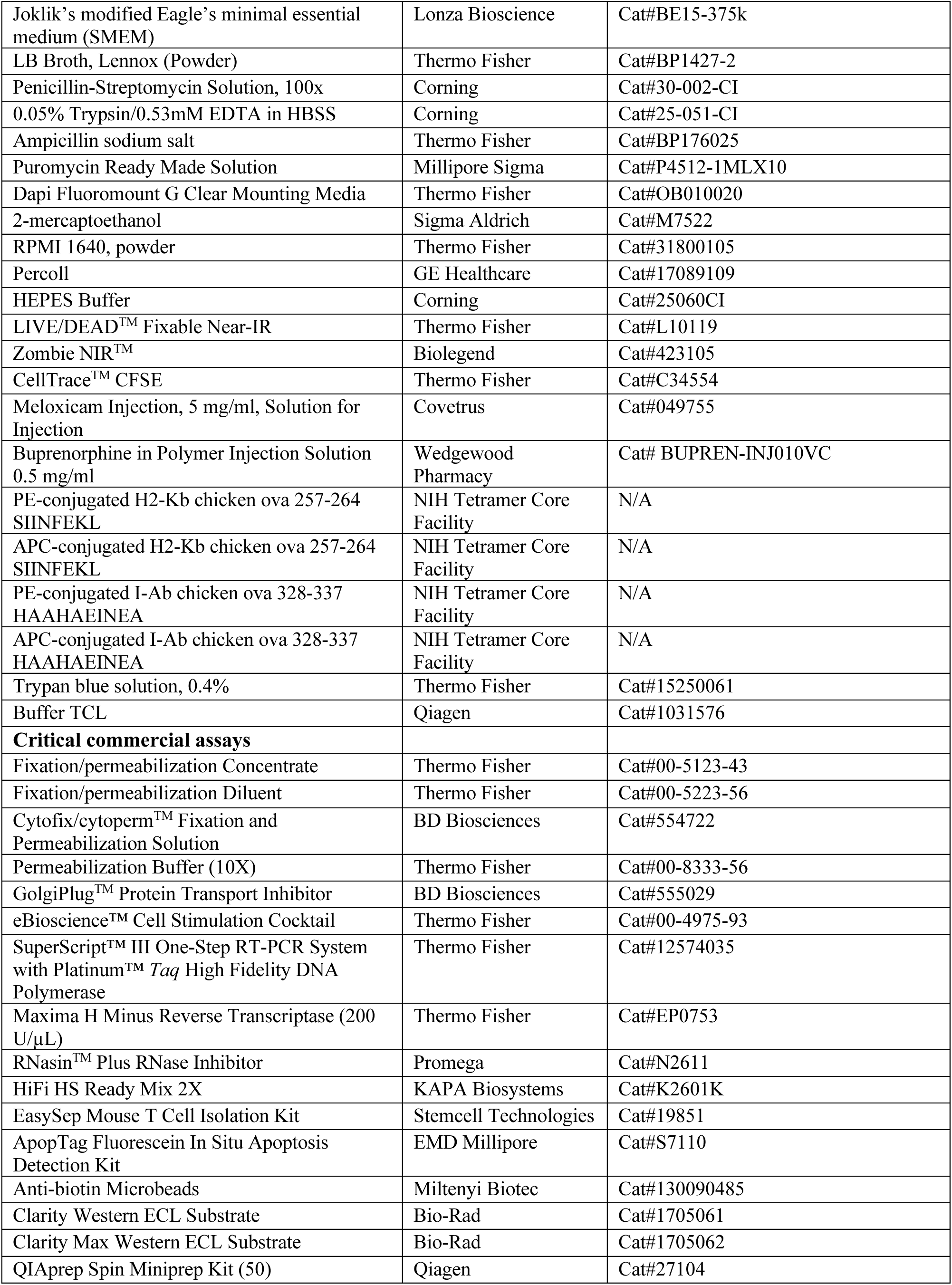
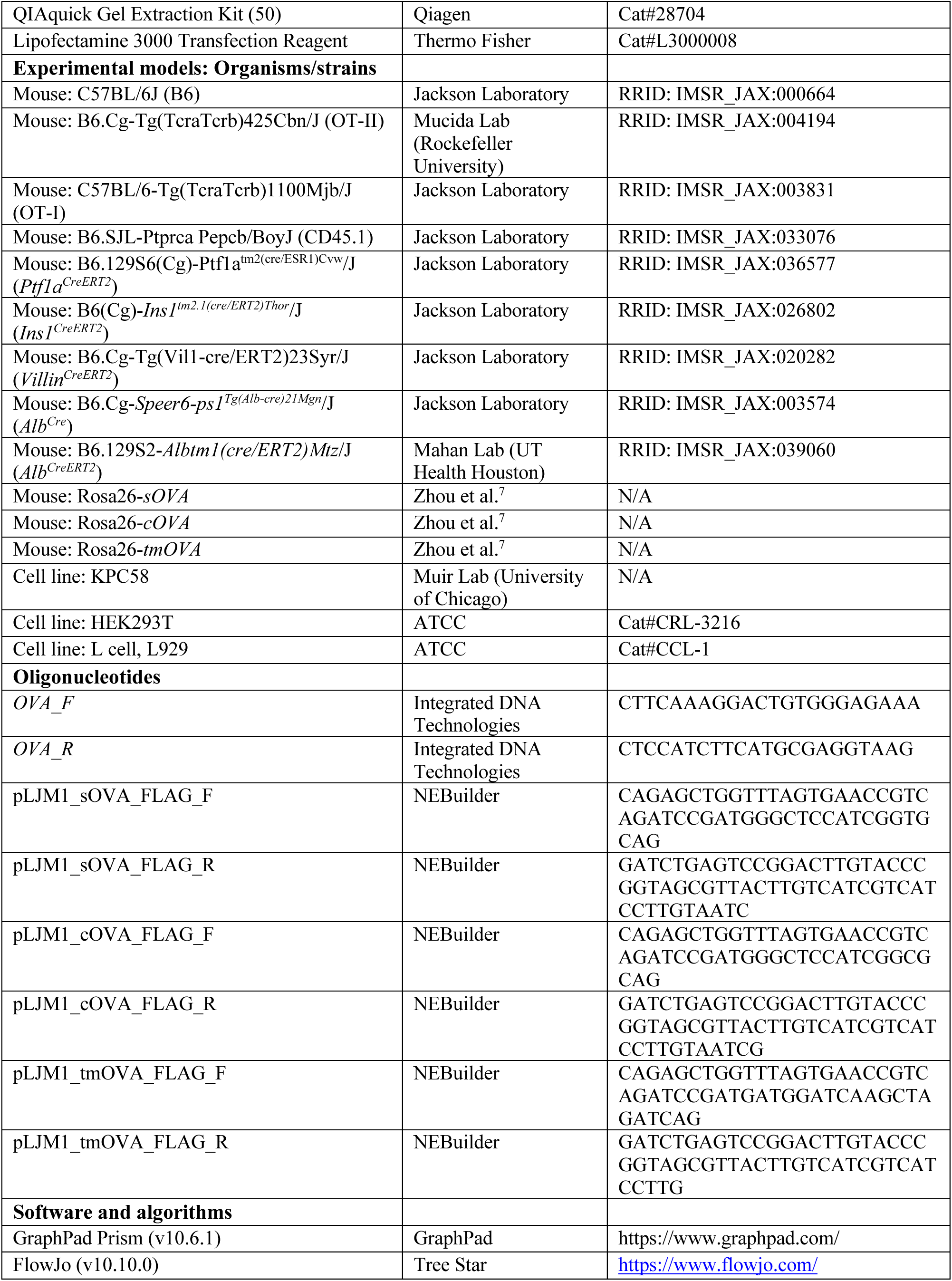
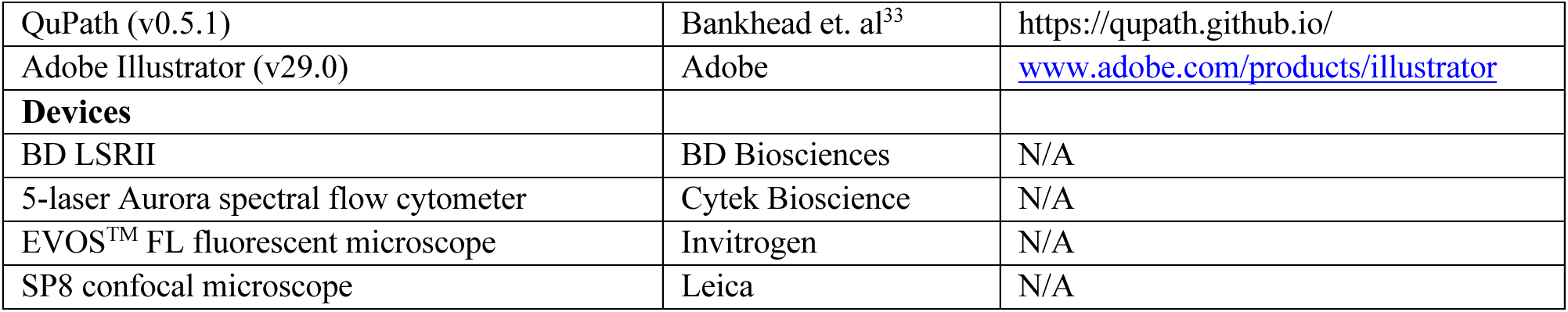

#### Mice

C57BL/6J, CD45.1 congenic (B6.SJL-Ptprc^a^ Pepc^b^/BoyJ), *Villin^CRE-ERT2^* (B6.Cg-Tg(Vil1-cre/ERT2)23 Syr/J), *Ptf1a^CRE-ERT2^*(B6.129S6(Cg)-Ptf1a^tm2^(cre/ESR1)Cvw/J), *Ins1^CRE-ERT2^*(B6(Cg)-Ins1^tm2.1(cre/ERT2)Thor^/J), OT-I (C57BL/6-Tg(TcraTcrb) 1100Mjb/J), OT-II (B6.Cg-Tg(TcraTcrb)425Cbn/J), *Alb*^Cre^ (B6.Cg-Speer6-ps1^Tg^(Alb-cre)21 Mgn/J), *p53*^LOXP^ (B6.129P2-*Trp53*^tm1Brn^/J) and *Kras*^FSF-G12D^ (B6(Cg)-*Kras*^tm5Tyj^/J) were purchased from The Jackson Laboratory. *Alb*^CreERT2^ mice were gifted by Dr. Kristin E. Mahan (UT Health Houston) after obtaining an MTA from the original investigator (Dr. Pierre Chambon, IGBMC, Illkirch, France) and rederived at our facility. The 3 lines of OVA mice were engineered in house and characterized in Zhou et, al^7^. All mice were maintained at the University of Chicago animal facility adhering to the AICUC protocol under specific pathogen-free (SPF) condition, confirmed to be free of segmented filamentous bacteria (SFB). Mice were used between 5-10 weeks of age. With the exception of *S. venezuelensis* infections (male mice only), a mix of male and female mice was used, with equal numbers of males and females in each experimental group.

#### Immunofluorescent staining of paraffin embedded slides

Tissues were fixed in 4% PFA/PBS for 2 h at RT and subjected to paraffinization. Paraffin-embedded tissue sections (5μm) were heated at 65°C for 20min, deparaffinized twice in xylene (10 min each), and rehydrated through a graded ethanol series. Sections were blocked for 30 min at RT with blocking buffer (5% donkey serum, 2.5% BSA in 1X PBS). Slides were permeabilized with 0.1% Triton X-100 before overnight incubation at 4°C in a humidified chamber with primary antibodies (see Antibodies section). The following day, slides were washed three times with PBST (PBS with 0.05% Tween-20) for 15 min each, incubated with appropriate secondary antibodies for 1 h at RT, washed again three times in PBST, and mounted with DAPI Fluoromount G Clear Mounting Media (SouthernBiotech). Images were acquired using SP8 Lightning confocal microscope (Leica) or Olympus VS200 Slideview Research Slide Scanner at the University of Chicago Microscopy Core.

#### Immunofluorescent staining of fresh frozen slides

Tissue was directly put into OCT compound and fresh frozen using pre-cooled 2-methylbutane bath and cut into 6-μm-thick sections. The slides were returned to RT right before staining and fixed in 1% PFA for 30 min. They were then blocked in 5% BSA + 0.1% Tween-20 in PBS for 1 hour at RT and incubated in primary antibodies diluted in blocking buffer overnight at 4°C. The slides were then washed, incubated with secondary antibodies (1:500 dilution) and mounted with DAPI Fluoromount G Clear Mounting Media. Stained slides were sent for whole slide scanning using Olympus VS200 Slideview Research Slide Scanner. Alternatively, pictures of slides were taken using SP8 Lightning Confocal Microscope (Leica).

#### Immunoblotting

Whole-cell or whole-tissue lysates were prepared in RIPA lysis buffer (20 mM HEPES, 0.2 mM EDTA, 300 mM NaCl, 1.5 mM MgCl_2_, 1% Triton X-100) supplemented with protease inhibitor (Roche) and 1 mM phenylmethylsulphonyl fluoride (PMSF; Sigma). Samples were separated by SDS-PAGE and transferred onto nitrocellulose membranes (Millipore). Membranes were probed with the indicated primary and secondary antibodies (see Key Resource Table).

#### Cell culture

HEK293T cells were purchased from ATCC and were cultured in Dulbecco’s Modified Eagle Media (DMEM; Gibco) supplemented with 10% FBS and 100 U/mL penicillin/streptomycin. KPC58 cell line was generously provided by Dr. Alexander Muir (University of Chicago)^16^. KPC cells were grown in RPMI-1640 media supplemented with 10% FBS and 100 U/mL penicillin and streptomycin. Spinner-adapted murine L929 cells for T1L titer determination were maintained in either suspension or monolayer cultures in Joklik’s modified Eagle’s minimal essential medium (SMEM; Lonza) supplemented to contain 5% fetal bovine serum (FBS; Gibco), 2 mM L-glutamine, 100 U/ml of penicillin, 100 μg/ml of streptomycin (Gibco), and 25 ng/ml of amphotericin B (Sigma). Cultures were maintained at 37°C in a humidified incubator with 5% CO_2_.

#### Lentiviral transduction of ovalbumin expression

pLJM1-Empty (Addgene #91980) plasmid was used as lentiviral vector, which was linearized by Nhel-HF (NEB) at 37°C for 20min and gel purified. Primers were designed using NEBuilder (https://nebuilder.neb.com/#!/) to create the necessary overlap with the plasmid. Insert fragments were created using the designed primers (listed in Key Resource Table) and Phusion High-fidelity DNA polymerase (NEB). The fragments were gel purified. Gibson assembly reaction was set up and the molar ratio for insert-to-vector was set to be 1:1. The reactants were incubated at 50°C for 2 h and the product was transformed into NEB Stable Competent E. coli cells (NEB #C3040H) according to the manufacturers’ protocol. Transformed E. coli cells were plated onto selection agar plates and single colonies were picked and sequences. The correct colonies were expanded, and plasmids were extracted using QIAGEN Plasmid Mini Kit.

Lentivirus was produced in HEK293T cells transfected at 80-90% confluency using Lipofectamine 3000 (Invitrogen) as recommended by the manufacturer and psPAX2 (Addgene) and pMD2.G (Addgene) packaging vectors. Medium was changed 8-12 hours after transfection and supernatant was collected after 48-72 h. Viral media was passed through a 0.45 μm filter and mixed with 10 μg ml^-1^ Polybrene (Sigma Aldrich) before being added to KPC cells. Infected cells were treated with puromycin to generate stable cell populations. OVA expression was verified by immunoblotting.

#### Flow Cytometry

Single-cell suspensions were first stained with LIVE/DEAD™ Fixable Near-IR Dead Cells Stain kits (Invitrogen) for 10 min at 4°C, followed by surface staining with antibody cocktails (1:200) for 20 min at 4°C. For intracellular cytokine detection, cells were fixed and permeabilized with Cytofix/cytoperm™ solution (BD Biosciences) for 20 min prior to staining with intracellular antibodies overnight at 4°C. For transcription factor analysis, cells were incubated in Fixation/permeabilization solution (eBiosciences) for 30 min to 2 hr, and subsequently stained with nuclear antibodies overnight at 4°C. All intracellular and nuclear antibodies were diluted 1:100 in 1x Permeabilization buffer (eBiosciences). Samples were acquired on an LSR II (BD Biosciences) or a 5-laser Aurora spectral flow cytometer (Cytek) and analyzed using FlowJo software. Cell division index was calculated using FlowJo formula as described (http://www.flowjo.com/v765/en/proliferation.html), whereby the index represents the fraction of total cell divisions normalized to the estimated starting cell number.

#### OVA tetramer staining

Single-cell suspensions were enriched for T cells using EasySep^TM^ Mouse T Cell Isolation Kit (Stemcell Technologies) and then incubated with Fc block for 10 min at 4°C to prevent nonspecific binding. For detection of OVA-specific CD8^+^ T cells, H-2Kb:OVA (SIINFEKL) PE- and APC-conjugated tetramers (NIH Tetramer Core Facility) were added at 1:200 in FACS buffer and incubated for 20 min at RT in the dark. For detection of OVA-specific CD4^+^ T cells, I-Ab:OVA (HAAHAEINEA) PE- or APC-conjugated tetramers (NIH Tetramer Core Facility) were added at 1:100 in T cell medium and incubated for 2 h at RT in the dark. Cells were then washed with FACS buffer before staining with surface antibody cocktail. Each tetramer lot was validated using OVA-immunized mice before use. Tetramer-positive gates were defined using OVA-naïve mice and fluorescence-minus-one controls.

#### Quantitative Real-time PCR

Total RNA from samples was extracted using TRIzol (Thermo Fisher Scientific), treated with DNase I (Invitrogen), and reverse transcribed into cDNA using SuperScript IV kit (Thermo Fisher Scientific) according to the manufacturer’s protocol. Quantitative real-time PCR was performed using Power SYBR Green PCR Master Mix (Applied Biosystems) on Quanstudio 6 Flex machine (Thermo Fisher Scientific). The relative expression of target genes was determined by normalization to housekeeping gene *36B4* and calculated using the formula of 2^−ΔCt^ ×10000. All data were averaged from at least 2 replicates.

#### Adaptive T cell transfer and CFSE tracing

Naïve T cells were isolated from pooled LNs and spleens by negative selection using biotinylated antibodies against NK1.1, B220, CD11c, CD11b, TER119 and either CD8α or CD4, followed by anti-biotin MACS beads (Miltenyi Biotec). Purity of transgenic cells was confirmed by flow cytometry (CD45.1^+^Vβ5^+^TCRβ^+^CD4^+^ for OT-II cells or CD45.1^+^Vβ5^+^TCRβ^+^CD8α^+^ for OT-I cells). Cell viability was determined by Trypan blue staining.

For experiments probing T cell outcomes in LNs, OT-II cells were labeled with CFSE Cell Proliferation Kit (Life Technologies) by incubating cells for 3-5 mins at 37°C, followed by a wash in PBS. A total of 750,000 OT-II or OT-I cells was transferred by retro-orbital injections under isoflurane anesthesia, and LNs harvested 4 days post-transfer. For experiments probing T cell responses in the tissues, transgenic cells were not CFSE-labeled. On the day of transfer, 200,000 OT-II and 50,000 OT-I cells were injected retro-orbitally. Organs were procured at indicated timepoints.

#### Induction of ovalbumin self-antigen in the intestine by tamoxifen administration

Rosa26-*OVA* mice were crossed to mice expressing tissue-specific Cre. Offspring were administrated tamoxifen-containing diet (TAM diet, Envigo) for 7 consecutive days or by daily intraperitoneal injection of 1mg tamoxifen (TAM) (Sigma) dissolved in 100 μl corn oil for 5 consecutive days at 4-6 weeks of age to induce for OVA self-antigen expression in the tissue of interest. Following TAM administration, mice were returned to a normal Chow diet for 4-7 days before use in experiments.

#### Oral ovalbumin administration

OVA (Sigma) was diluted in sterile PBS and filter sterilized prior to administration. Mice received 25 mg OVA in 200 μl PBS by oral gavage one time, one day post OT-I/OT-II co-transfer.

#### Infections

Reovirus strain type 1 Lang was recovered using plasmid-based reverse genetics^34^ and purified using CsCl_2_ gradient centrifugation^35^. Titers of purified virus stocks were determined by plaque assay using L929 cells. Intragastric infection was performed by oral gavage mice with 1×10^9^ PFU T1L in 200 μl PBS^35^. Intranasal infection was performed by lightly anesthetizing mice with isoflurane and inoculating intranasally with 1 × 10^9 PFU of T1L diluted in PBS in a total volume of 50 μl (25 μl per nostril).

*S. venezuelensis* was maintained in NSG mice by subcutaneous infection with 10,000-30,000 larvae. Fecal pellets from infected NSG mice were collected and spread on Whatman paper, and incubated in water at 28°C. Larvae emerging over 3-4 days were collected to reinitiate the infection cycle or to be used for infection of experimental mice. For experimental mouse infection, 1000 larvae were administrated subcutaneously to male mice. Successful infection was confirmed by appearance of eggs in the fecal content of the infected mice. OT-II cells were transferred 5 days post infection.

Adeno-Cre virus was diluted in PBS to a final concentration of 5 × 10^5 PFU/ml. A total volume of 200 μl of the viral suspension was administered to mice via tail vein injection. OT-II cells were transferred 14 days post infection.

#### Carbon tetrachloride (CCl_4_) administration

CCl_4_ was diluted in corn oil at 20% (v/v). A 50 μl volume of this solution was administered intraperitoneally to a 20 g mouse, delivering a dose of 50 μl per kg body weight.

#### Concanavalin A (Con A) administration

Con A was reconstituted from lyophilized powder into a 2.4 mg/mL working stock using sterile PBS. 100 μl was administered retro-orbitally to a 20 g mouse, delivering a dose of 12 mg/kg.

#### Cerulein administration

6 hourly intraperitoneal injections of 50ug/kg cerulein (Bachem) on each injection day while control mice were injected with the same amount of saline. 3 injection days was performed at 7, 4, 1 day before OT-II cell transfer.

#### Streptozotocin (STZ) administration

Prior to STZ treatment, mice were fasted for 4-5 hours. STZ was dissolved in sterile sodium citrate buffer immediately prior to i.p. injection. Sodium citrate buffer was prepared by adding 1.47 g sodium citrate to 50 mL H2O and adjusting pH to 4.5 with citric acid. Mice were given a single high dose (150 mg/kg).

#### ApopTag staining and apoptotic index

Apoptosis was detected using ApopTag Fluorescein In Situ Apoptosis Detection Kit (Sigma, S7110). Paraffin sections were deparaffinized and rehydrated as described for immunofluorescence staining, followed by ApopTag staining according to the manufacturer’s protocol. The stained slides were scanned and analyzed in QuPath. Apoptotic index represents the percentage of ApopTag^+^ cells over total cell number.

#### Isolation of T cells from lymph nodes

For T cell analysis, LNs were dissected into RPMI supplemented with 2% NCS and 1mM EDTA, mashed between two frosted glass slides, and filtered into 96-well plates. Cells were immediately subjected to Live/Dead and surface marker staining.

#### Isolation of T cells from the liver and pancreas

For hepatic immune cell isolation, liver was perfused with approximately 3ml of collagenase D and DNase I in RPMI through the portal vein. The liver was then digested at 37°C for 16-18 min. The reaction was quenched with ice-cold RPMI with 10% FBS. Parenchymal cells were discarded by centrifugation (75g for 1 min). The supernatant was centrifuged again at 800g for 2 min. The pellet was resuspended in 5 ml of Histopaque 1077 and gently overlayed with 10 ml of RPMI. Immune cells were enriched through gradient centrifugation at 900g for 18 min at RT with slow acceleration and deceleration.

For pancreatic immune cell isolation, pancreas was perfused with 2-3ml of collagenase IV and DNase I in HBSS with Ca and Mg through the bile duct. LNs were detached from the pancreas using forceps. The pancreas was then digested at 37°C for 25 min with repeated mechanical shacking every 5min throughout the digestion. The reaction was quenched with ice-cold RPMI with 10% FBS. The samples were filtered and centrifuged at 800g for 2 min. The pellet was resuspended in 5 ml of Histopaque 1077 and gently overlayed with 10 ml of RPMI. Immune cells were enriched through gradient centrifugation at 900g for 18 min at RT with slow acceleration and deceleration. Cells were immediately subjected to Live/Dead and surface marker staining.

#### T cell stimulation

For the detection of OT-I cytokine production, single cell suspension of isolated lymphocytes was pelleted by centrifugation and resuspended in T cell media containing cell stimulation cocktail (1:500; eBioscience). Cultures were incubated for 4 h at 37°C. Following stimulation, cells were pelleted, washed once with fresh medium, and processed for cell staining.

#### Pancreatic cancer induction

Ptf1a^CreERT2/w^*Trp53*^fl/fl^Kras^LSL-G12D/w^sOVA^+/w^ (PDAC-sOVA) mice were fed with tamoxifen-containing diet (Envigo # TD.130858) for 7 days at 3-4 weeks of age for induction of tumorigenesis and OVA expression. OT-I and OT-II cells were transfer at day 7 post induction. For T cell analysis in the LNs, LNs were harvested at day 10 post termination of the tamoxifen diet. For tumor area analysis, pancreas was harvested at day 28 post induction.

#### Orthotopic model for pancreatic cancer

Mice were treated with Meloxicam (0.5 mg) and Buprenorphine (10 μg per mouse) prior to the surgery for pain management. Mice were then anesthetized using isoflurane. 5k or 50k of KPC cells expressing sOVA, cOVA or empty vector control were surgically implanted to the mid portion of the pancreas. Surgical incision was sutured, and mice were monitored for the need of further pain management. At day 11 post operation, OT-I and OT-II cells were co-transferred and mice were infected with T1L or vehicle control. For T cell analysis in the LNs, LNs were harvested at day 14 post operation. For tumor analysis, mice were sacrificed at day 22-23 post operation and tumor size / weight was measured. Tumor size was calculated as 0.5 × length × width^2^. Tumors were fresh frozen in OCT for downstream immunofluorescent analysis.

#### Pancreatic lesion quantification

Whole-pancreas H&E sections, including tumors, were scanned and analyzed using QuPath. Spleen and lymph node regions were excluded, and the total pancreatic area was outlined. Each lesion was annotated, and lesion number and total lesion area were recorded. The percentage of lesion area was calculated relative to the total pancreatic area.

#### Administration of CD8b neutralizing antibody

One week after T1L infection, mice received 100 μg of CD8β-neutralizing antibody (SelleckChem) per mouse via intraperitoneal injection every 5 days. Control mice received 100 μg of isotype control antibody on the same schedule.

## Statistical analysis

All data were analyzed with Prism software (GraphPad). Data is presented as average ± SEM. Comparison between two treatment conditions by two-tailed unpaired Student’s t-test assuming a Gaussian distribution. Gaussian distribution tests appropriate for small sample sizes were applied for all datasets (Shapiro-Wilk and Kolmogorov-Smirnov test); all data sets passed the requirements for Gaussian distribution. P-values or representative symbols are noted when differences are less than or equal to 0.1; all other differences were not found to be significant (n.s., p > 0.1). ^∗^p < 0.05, ^∗∗^p < 0.01, ^∗∗∗^ p < 0.001). One-way ANOVA was used for multivariant comparisons between more than two groups. Two-tailed t-test was used to compare two groups (i.e. NI vs. infected LNs).

---

**Figure S1.**
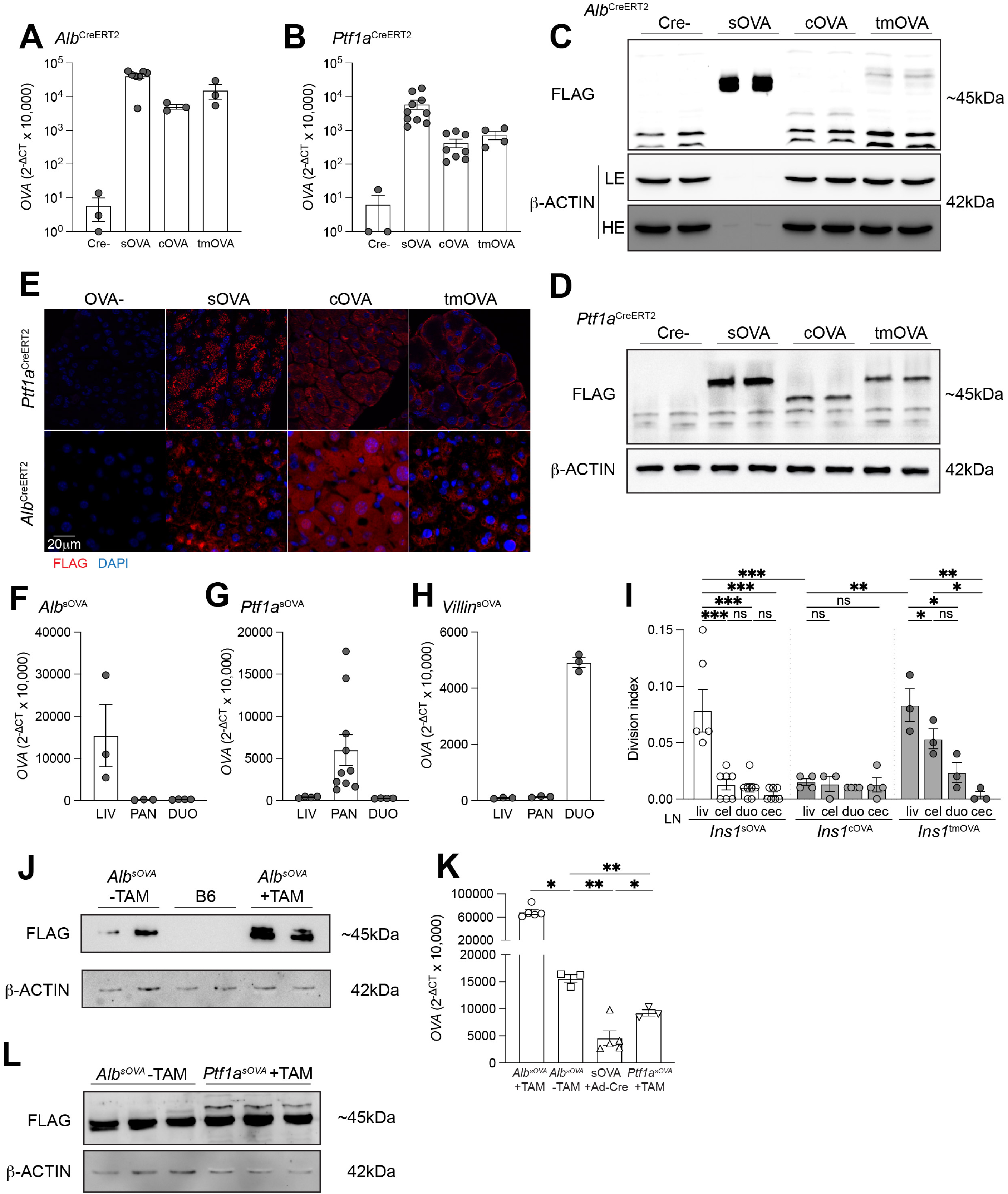
Tissue-specific tolerance mechanism, subcellular source of self-antigen and antigen dose dictate T cell fate of self-reactive CD4^+^ T cells at homeostasis. **A-B.** Expression levels of *OVA* in the liver (**A**) and pancreas (**B**) of mice with indicated genotypes (*n* = 3-10 per group, as indicated by symbols). **C-D**. Hepatic (**C**) and pancreatic (**D**) tissue lysates from mice with indicated OVA form and their CRE-negative controls were immunoblotted using antibodies against FLAG, and β-ACTIN as a loading control. **E**. Sections of pancreas (top) and liver (bottom) from mice with indicated genotypes stained for FLAG and DAPI (nuclei). Scale bar represents 20μm. F-H. *OVA* expression levels in the liver, pancreas and duodenum tissue of mice with indicated genotypes, showing tissue-specific expression restricted to the intended sites (*n* = 3-10 per group, as indicated by symbols). **I.** Division index of OT-II cells in the indicated LNs 6 days after transfer into *Ins1^sOVA^*, *Ins1^cOVA^* and *Ins1^tmOVA^* mice (*n* = 4-7 per group, as indicated by symbols). **J**. Hepatic tissue lysates from *Alb^sOVA^* mice treated with or without tamoxifen and B6 mice were immunoblotted using antibodies against FLAG and β-ACTIN. K. *OVA* expression levels in liver tissue of *Alb^sOVA^* mice treated with or without tamoxifen and sOVA mice infected with adeno-Cre virus and pancreas of *Ptf1a^sOVA^* mice (*n* = 3-5 per group, as indicated by symbols). **L**. Hepatic tissue lysates from *Alb^sOVA^*mice without tamoxifen treatment and pancreatic lysates from tamoxifen-treated *Ptf1a^sOVA^* mice were immunoblotted using antibodies against FLAG and β-ACTIN. Data are pooled from two independent experiments in A, B, F-I. * p<0.05, ** p<0.01, *** p<0.001 by t-test.

**Figure S2.**
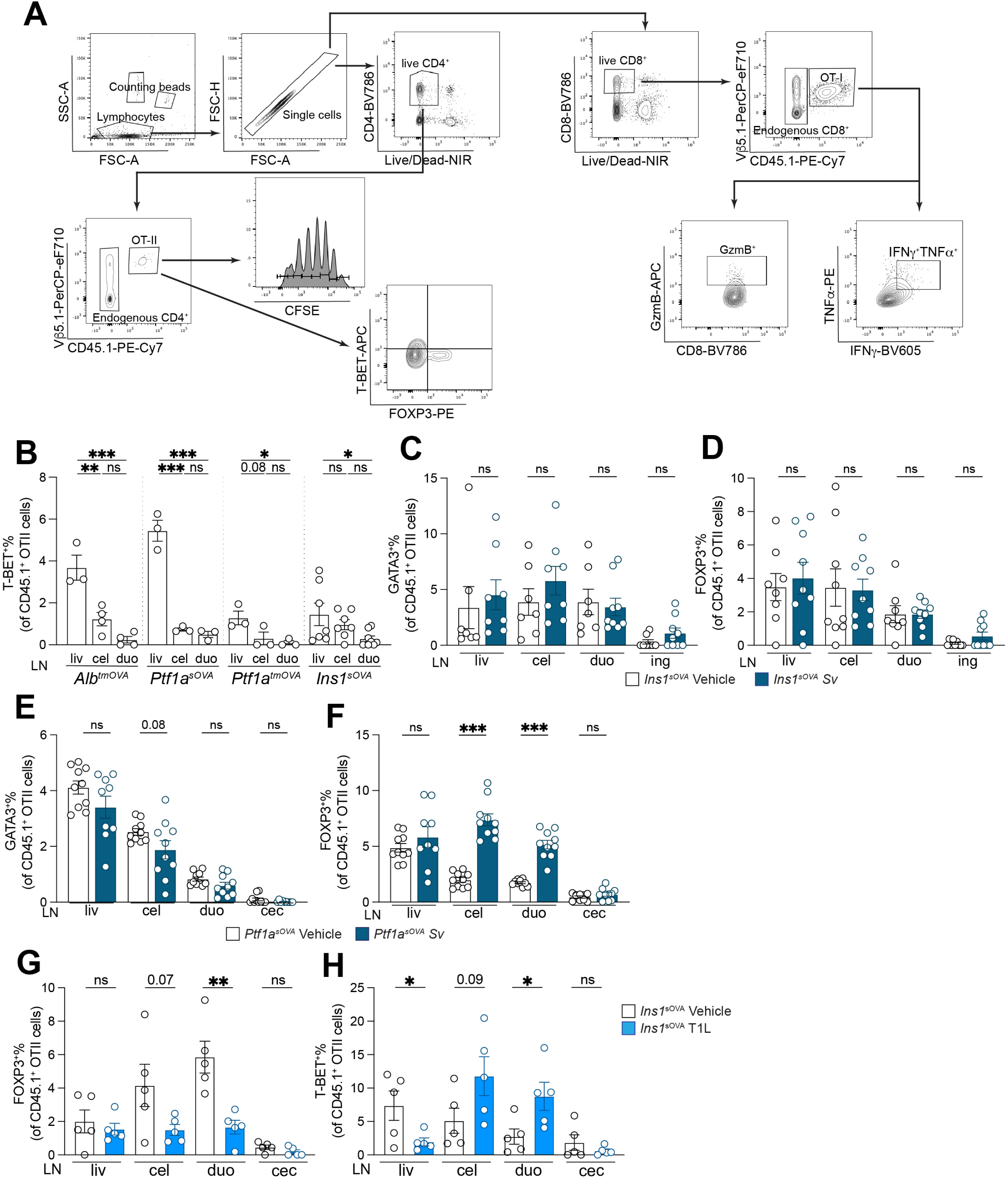
Type 1, but not type 2, intestinal infection renders pancreas-reactive T cells pro-inflammatory. **A.** Gating strategy for OT-II proliferation and cell fate of both OT-I and OT-II cells. **B.** Frequencies of T-BET^+^ among total OT-II cells in the indicated LNs 96 h after transfer into *Alb^tmOVA^*, *Ptf1a^sOVA^*, *Ptf1a^tmOVA^* and *Ins1^sOVA^* mice (*n* = 3-8 per group, as indicated by symbols). **C-F.** Frequencies of GATA3^+^ (**C, E**) or FOXP3^+^ (**D, F**) among total OT-II cells 72 h after transfer into *Ins1^sOVA^* (**C-D**) or *Ptf1a^sOVA^* (**E-F**) mice that were *S. venezuelensis*- or mock-infected (*n* = 8-10 per group, as indicated by symbols). **G-H.** Frequencies of FOXP3^+^ (**G**) or T-BET^+^ (**H**) among total OT-II cells in indicated LNs 72 h after transfer into T1L- or mock-infected *Ins1^sOVA^* mice (*n* = 5 per group). Data are pooled from two independent experiments in B-H. * p<0.05, ** p<0.01, *** p<0.001 by t-test.

**Figure S3.**
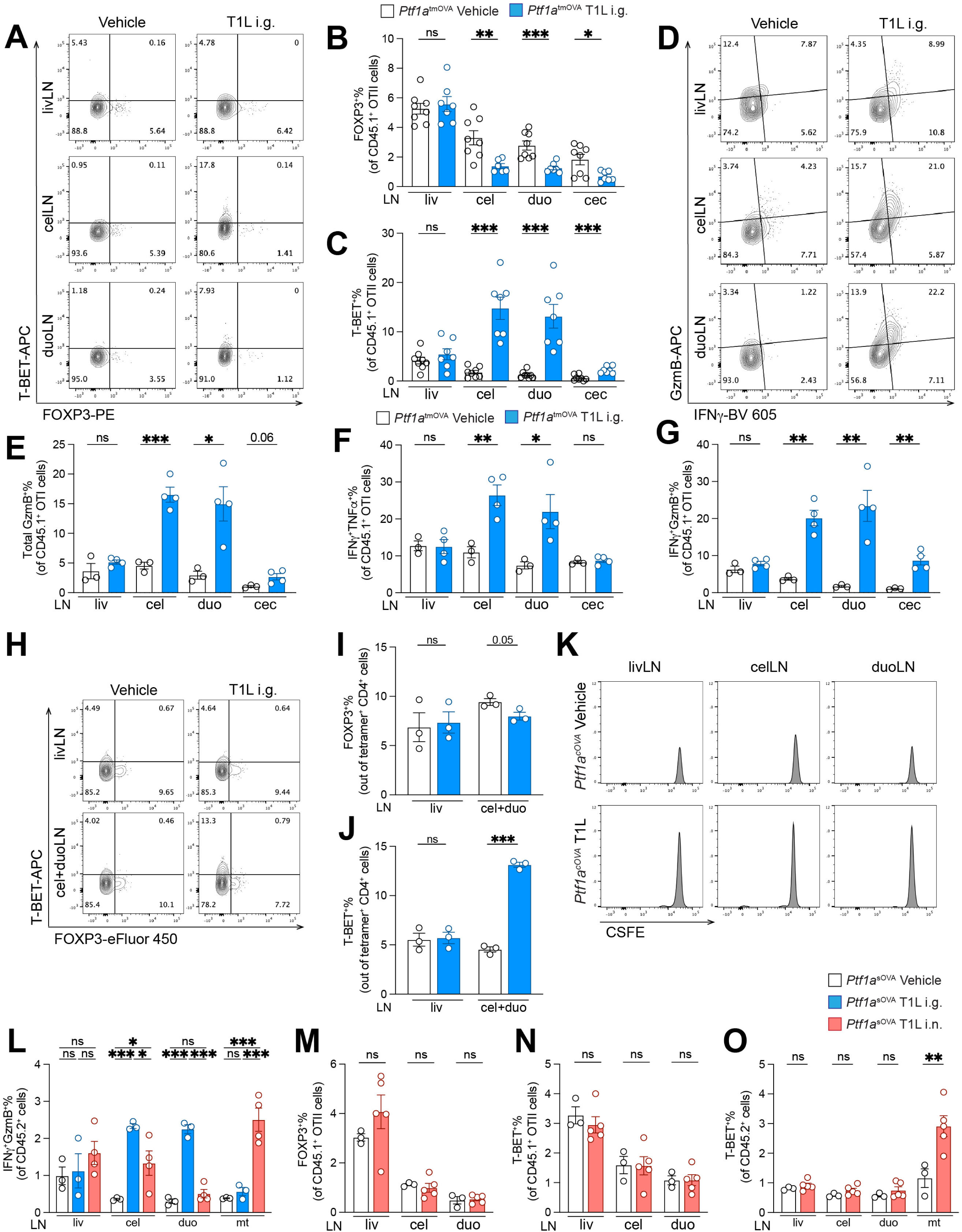
Intestinal reovirus infection renders pancreas-reactive T cells pro-inflammatory by LN co-drainage. **A.** Plots of OT-II cells in LNs of *Ptf1a^tmOVA^* mice that are inoculated perorally with T1L or vehicle control, representative gating applied in (**B**) and (**C**). **B-C.** Frequencies of FOXP3^+^ (**B**) or T-BET^+^ (**C**) among total OT-II cells in indicated LNs 72 h after transfer into mice in (**A**) (*n* = 7-8 per group, as indicated by symbols). **D.** Plots of OT-I cells in LNs of *Ptf1a^tmOVA^* mice that are inoculated perorally with T1L or vehicle control. **E-G.** Frequencies of total GzmB^+^ (**E**), IFNγ^+^GzmB^+^ (**F**) or IFNγ^+^TNFα^+^ (**G**) among total OT-I cells in indicated LNs 72 h after transfer into mice in (**D**) (*n* = 3-4 per group, as indicated by symbols). **H-J**. Flow plots (**H**) and frequencies of FOX3^+^ (**I**) or T-BET^+^ (**J**) among total OVA-tetramer^+^CD4^+^ T cells in LNs of T1L-or vehicle-infected *Ptf1a^sOVA^* mice (*n* = 3 per group). **K.** CFSE flow plots of OT-II cells in indicated LNs from *Ptf1a^cOVA^*mice orally infected with T1L or given vehicle control. **L.** Frequencies of endogenous IFNγ^+^GzmB^+^ among total CD8^+^ cells in LNs of *Ptf1a^sOVA^* mice that are either intragastrically or intranasally infected with T1L (*n* = 3-4 per group, as indicated by symbols). **M-O.** Frequencies of FOXP3^+^ (**M**), T-BET^+^ (**N**) among total OT-II cells or endogenous T-BET^+^ - among total CD4^+^ cells (**O**) in indicated LNs of T1L-infected or vehicle-infected *Ptf1a^sOVA^* mice (*n* = 3-5 per group, as indicated by symbols). Data are pooled from two independent experiments in B, C. Data represent one experiment in E-G, I, J, L-O. * p<0.05, ** p<0.01, *** p<0.001 by t-test.

**Figure S4.**
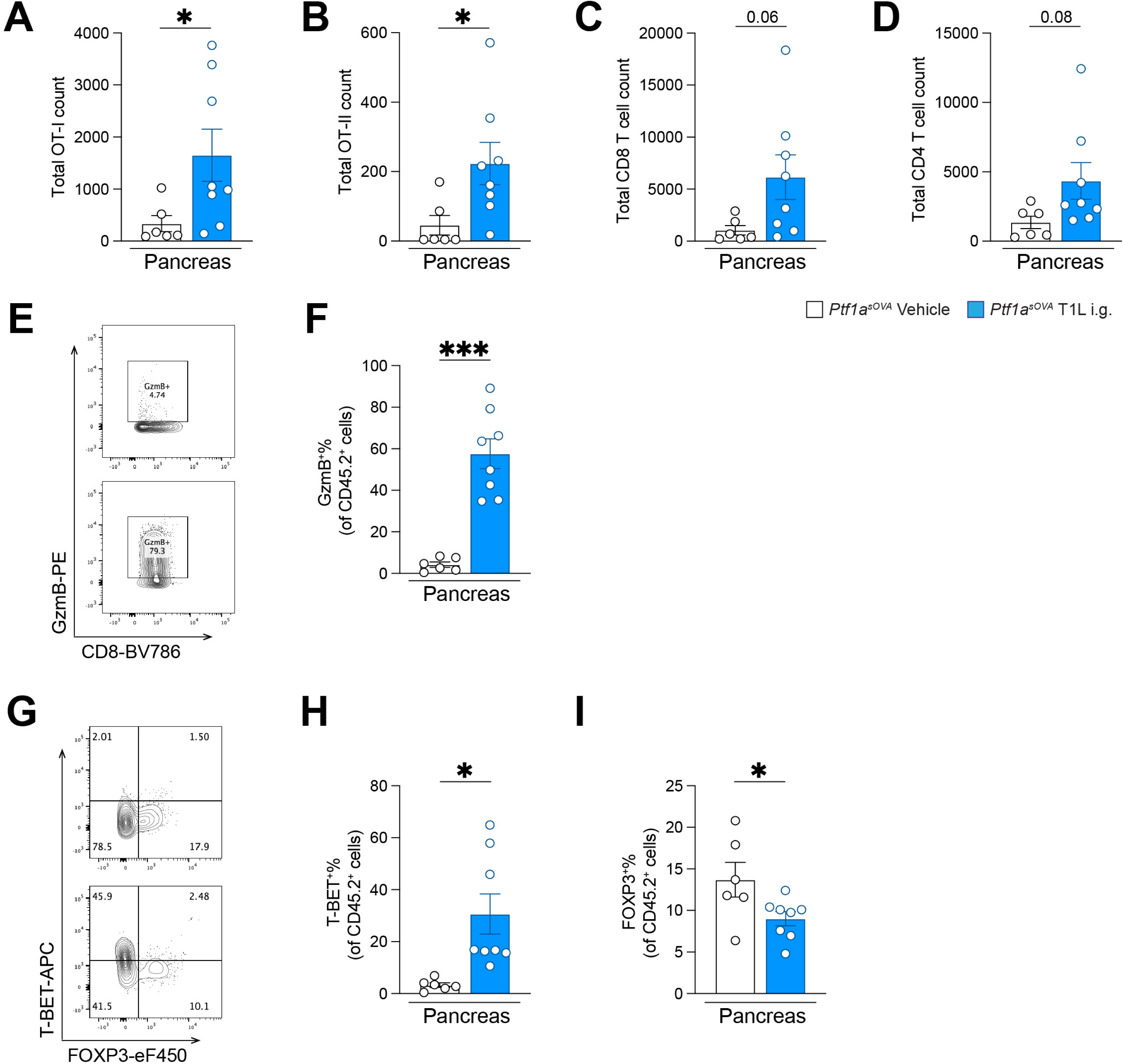
Intestinal reovirus-induced pro-inflammatory T cells infiltrate the pancreas and cause severe tissue destruction. **A-D.** Total OT-I (**A**), OT-II (**B**), CD8^+^ (**C**) and CD4^+^ (**D**) T cell numbers in the pancreas of orally T1L-infected or vehicle-infected *Ptf1a^sOVA^*mice 7 days post transfer. **E-F.** Representative plots of endogenous CD8^+^ T cells (**E**) and frequencies of GzmB^+^ among total CD8^+^ T cells (**F**) in the pancreas of T1L-infected or vehicle-infected *Ptf1a^sOVA^*mice. **G-I.** Representative flow plots of endogenous CD4^+^ T cells (**G**) and frequencies of T-BET^+^ (**H**) or FOXP3^+^ among total CD4^+^ T cells (**I**) in the pancreas of T1L-infected or vehicle-infected *Ptf1a^sOVA^* mice. Data are pooled from two independent experiments (*n* = 6-8 per group, as indicated by symbols). * p<0.05, ** p<0.01, *** p<0.001 by t-test.

**Figure S5.**
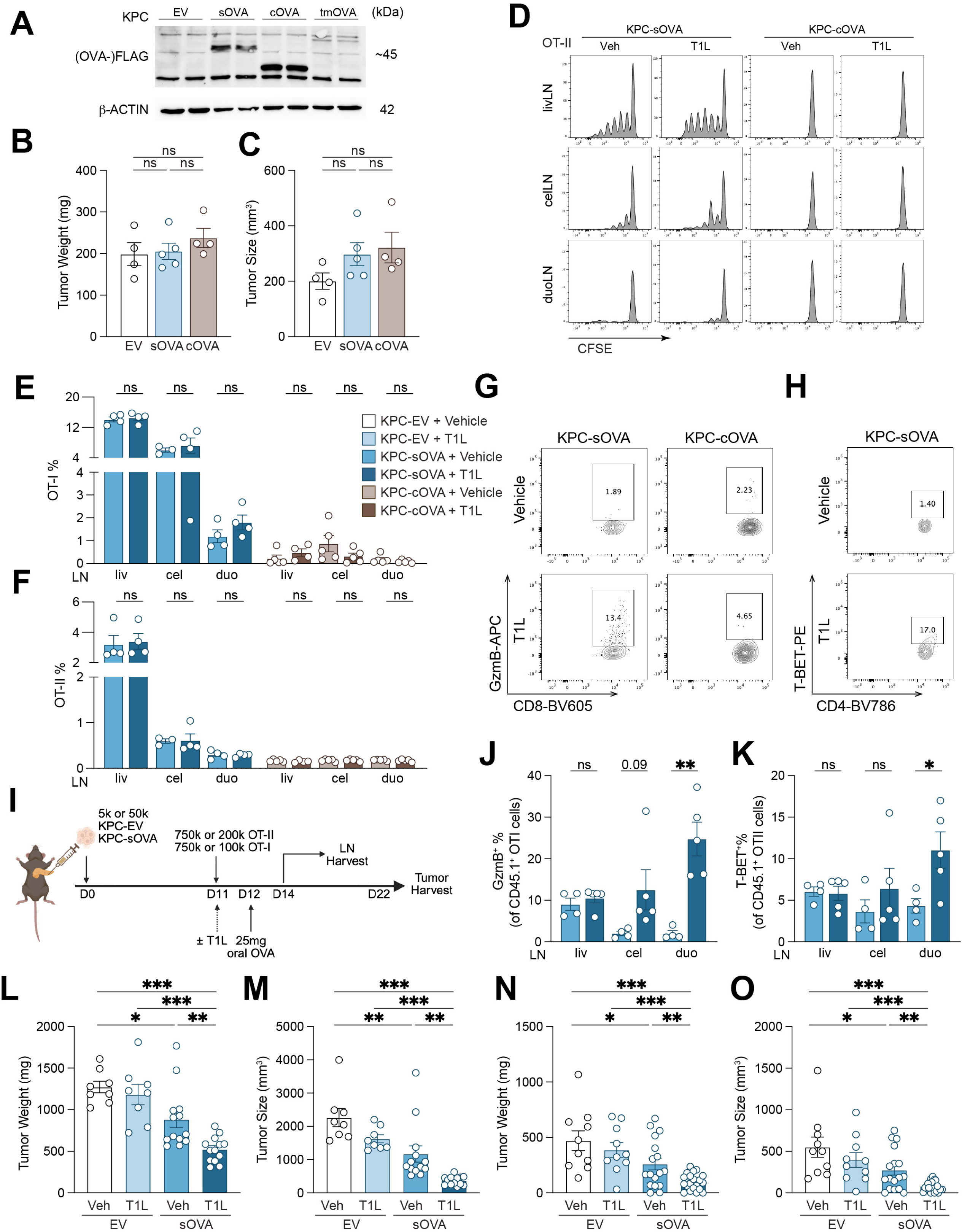
Intestinal reovirus infection enhances T cell infiltration and tumor control in an orthotopic PDAC model. **A.** Western blot of cell lysates of KPC cells expressing indicated OVA forms or empty vector control using antibodies against FLAG and β-ACTIN. **B-C.** Tumor weights (**B**) and sizes (**C**) at day 11 post KPC cell injection were measured to confirm that OVA expression does not alter tumor growth (*n* = 4-5 per group, as indicated by symbols). **D.** CFSE dilution flow plots of OT-II cells in the indicated LNs from orally T1L-infected or mock-infected mice receiving sOVA- or cOVA-expressing KPC tumor cells. **E-F**. Frequencies of OT-I among total CD8^+^ cells (**E**) or OT-II among total CD4^+^ (**F**) cells in indicated LNs of mice receiving sOVA- or cOVA-expressing KPC tumor cells (*n* = 4-5 per group, as indicated by symbols). **G-H**. Plots of OT-I cells (**G**) or OT-II (**H**) cells in duoLNs of mice receiving sOVA- or cOVA-expressing tumor cells. **I**. Schematic of orthotopic PDAC model for assessing T cell fate in the LNs and tumor progression. **J-K**. Frequencies of GzmB^+^ among total OT-I cells (**J**) or T-BET^+^ among total OT-II (**K**) cells in indicated LNs of mice receiving 50k sOVA-expressing KPC tumor cells with one dose of OVA gavage 1 day post cell transfer (*n* = 4-5 per group, as indicated by symbols). **L-M.** Weights (**L**) and sizes (**M**) of tumors from mice receiving 50k indicated tumor cells and one dose of OVA gavage 1 day post cell transfer (*n* = 8-13 per group). **N-O.** Weights (**N**) and sizes (**O**) of tumors from mice receiving 5k indicated tumor cells and one dose of OVA gavage 1 day post cell transfer (*n* = 10-20 per group, as indicated by symbols). Data are pooled from two independent experiments in L-O. Data represent one experiment in B, C, E, F, J, K. * p<0.05, ** p<0.01, *** p<0.001 by t-test.

**Figure S6.**
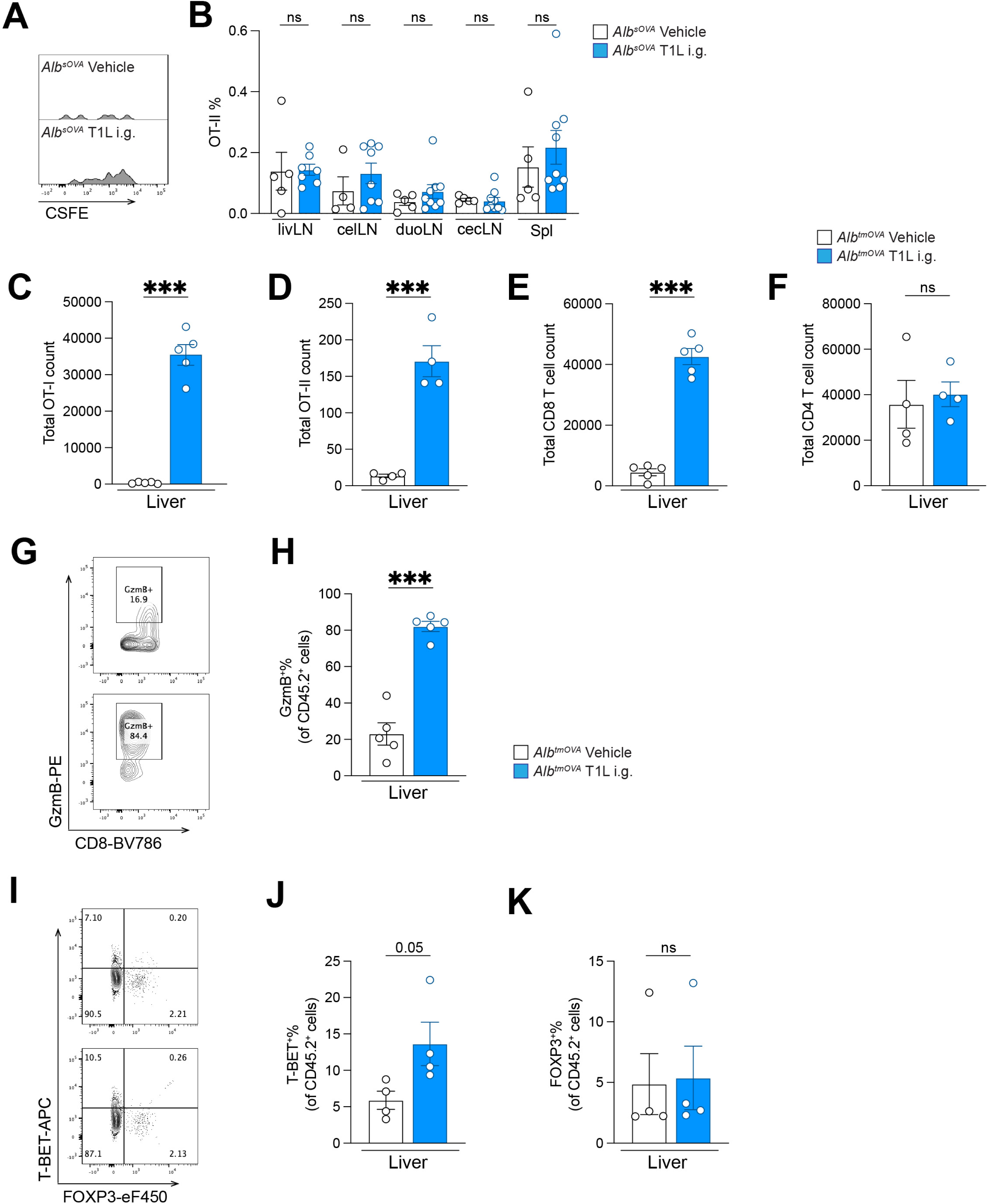
Intestinal reovirus infection induces liver-reactive T cells and hepatic damage by LN co-drainage. **A-B.** CFSE flow plots (**A**) and frequency of OT-II among total CD4^+^ cells (**B**) in indicated LNs and spleen from *Alb^sOVA^* mice orally infected with T1L or treated with vehicle control (*n* = 5-9 per group, as indicated by symbols). **C-F.** Total OT-I (**C**), OT-II (**D**), CD8^+^ (**E**) and CD4^+^ (**F**) T cell numbers in the liver of T1L- or vehicle-infected *Alb^tmOVA^* mice 7 days post transfer. **G-H.** Representative plots of endogenous CD8^+^ T cells (**G**) and frequencies of GzmB^+^ among total CD8^+^ T cells (**H**) in the liver of T1L- or mock-infected *Alb^tmOVA^*mice. **I-K.** Representative plots of endogenous CD4^+^ T cells (**I**) and frequencies of T-BET^+^ (**J**) or FOXP3^+^ among total CD4^+^ T cells (**K**) in the liver of T1L- or mock-infected *Alb^tmOVA^* mice. Data are pooled from two independent experiments in A-B. Data are representative of two independent experiments (*n* = 4 per group). * p<0.05, ** p<0.01, *** p<0.001 by t-test.

**Figure S7.**
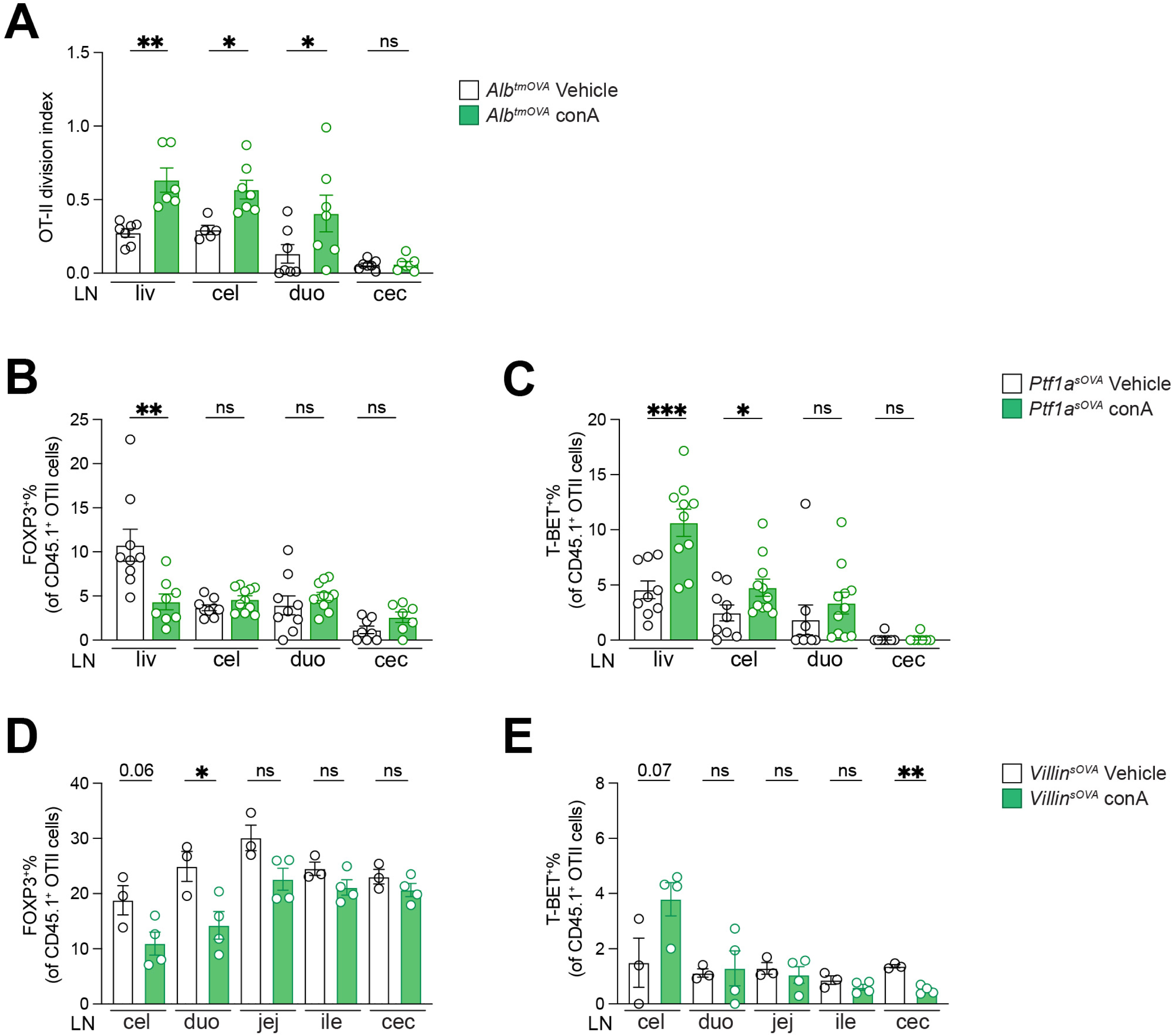
Con A-induced liver injury promotes pro-inflammatory conversion of pancreas-and gut-reactive T cells. **A.** Division index of OT-II cells in indicated LNs 96 h after transfer into con A- or vehicle-treated *Alb^tmOVA^* mice (*n* = 6 per group). **B-C.** Frequencies of FOXP3^+^ (**B**) or T-BET^+^ (**C**) among total OT-II cells in indicated LNs 96 h after transfer into con A- or vehicle-treated *Ptf1a^sOVA^* mice (*n* = 9-10 per group, as indicated by symbols). **D-E.** Frequencies of FOXP3^+^ (**B**) or T-BET^+^ (**C**) among total OT-II cells in indicated LNs 96 h after transfer into con A- or vehicle-treated *Villin^sOVA^*mice (*n* = 3-4 per group, as indicated by symbols). Data are pooled from two independent experiments in A-C. Data represent one experiment in D-E. * p<0.05, ** p<0.01, *** p<0.001 by t-test.

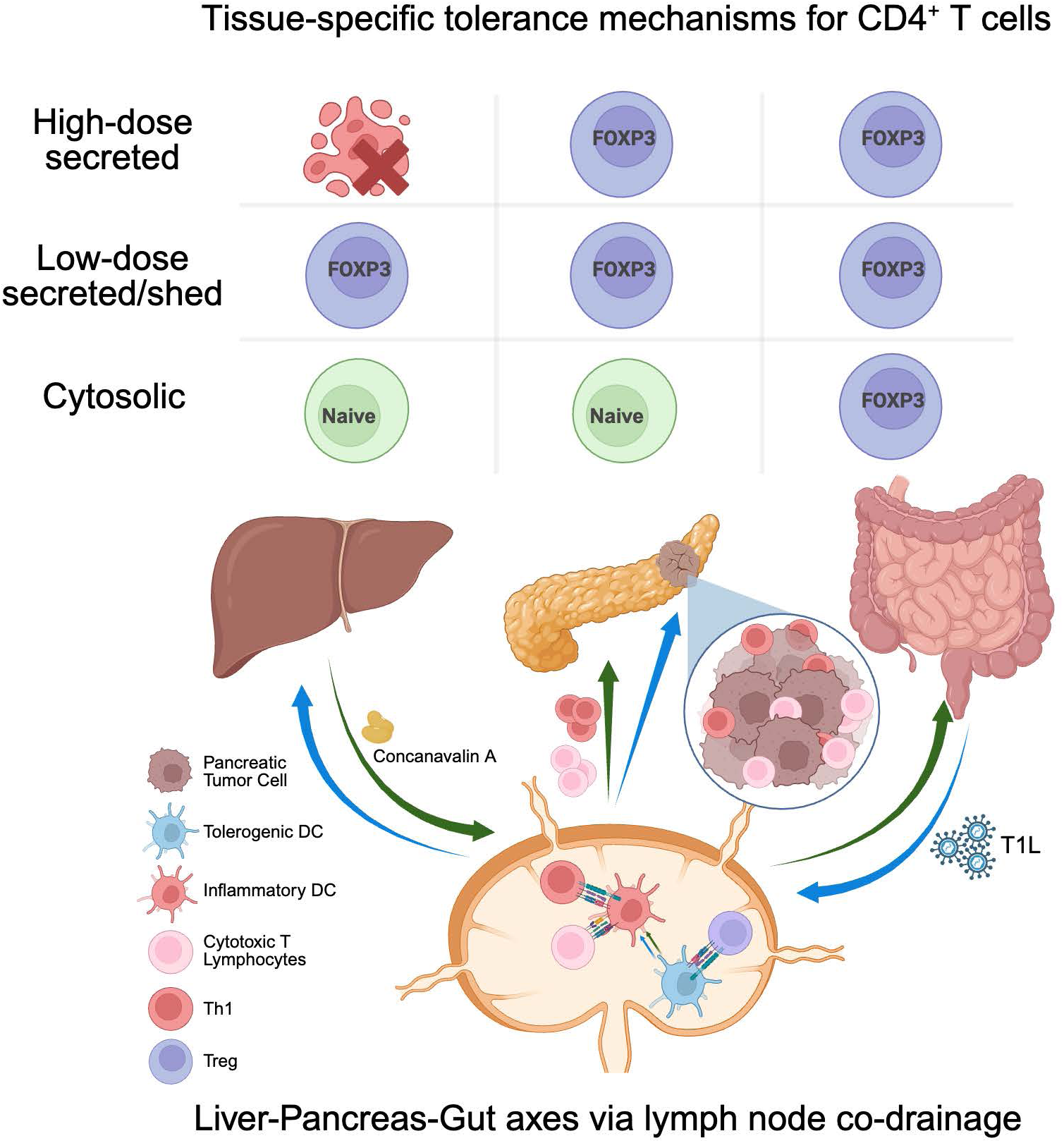

